# Asynchronous abundance fluctuations can drive giant genotype frequency fluctuations

**DOI:** 10.1101/2024.02.23.581776

**Authors:** Joao A Ascensao, Kristen Lok, Oskar Hallatschek

## Abstract

Large stochastic population abundance fluctuations are ubiquitous across the tree of life^1–7^, impacting the predictability of population dynamics and influencing eco-evolutionary outcomes. It has generally been thought that these large abundance fluctuations do not strongly impact evolution, as the relative frequencies of alleles in the population will be unaffected if the abundance of all alleles fluctuate in unison. However, we argue that large abundance fluctuations can lead to significant genotype frequency fluctuations if different genotypes within a population experience these fluctuations asynchronously. By serially diluting mixtures of two closely related *E. coli* strains, we show that such asynchrony can occur, leading to giant frequency fluctuations that far exceed expectations from models of genetic drift. We develop a flexible, effective model that explains the abundance fluctuations as arising from correlated offspring numbers between individuals, and the large frequency fluctuations result from even slight decoupling in offspring numbers between genotypes. This model accurately describes the observed abundance and frequency fluctuation scaling behaviors. Our findings suggest chaotic dynamics underpin these giant fluctuations, causing initially similar trajectories to diverge exponentially; subtle environmental changes can be magnified, leading to batch correlations in identical growth conditions. Furthermore, we present evidence that such decoupling noise is also present in mixed-genotype *S. cerevisiae* populations. We demonstrate that such decoupling noise can strongly influence evolutionary outcomes, in a manner distinct from genetic drift. Given the generic nature of asynchronous fluctuations, we anticipate that they are widespread in biological populations, significantly affecting evolutionary and ecological dynamics.

## Introduction

The dynamics of evolution fundamentally depends on the interplay between the deterministic and stochastic forces acting on populations. Natural selection pushes allele frequencies up or down in the population, depending on the relative allele fitness, while genetic drift causes random allele frequency fluctuations, with no preferred direction^8–12^. Theoretical population genetics has provided many examples of how natural selection and genetic drift can interact with each other. For example, the probability that a mutant will establish in a population is determined primarily by the stochastic dynamics dominated by genetic drift at low mutant frequencies, followed by deterministic dynamics dominated by natural selection at higher frequencies^10,13,14^. Even in purely neutral scenarios, stochastic forces alone can often lead to surprisingly complex evolutionary dynamics^15–18^.

Ecological dynamics can also be strongly influenced by stochastic demographic fluctuations. It has long been noted that populations across the tree of life exhibit strong abundance fluctuations, nearly universally^6,19,20^. In many of the documented cases, the abundance fluctuations follow Taylor’s power law^1–5^–a power-law relationship between the mean and variance of the abundance fluctuations. Many different ecological processes can cause abundance fluctuations that obey Taylor’s law, including fluctuating environments^21^, spatial effects^1,22^, or chaotic dynamics^23,24^. Chaotic population dynamics in particular have captured the interest of ecologists for decades, ever since it was noted that even simple models of single populations can display chaotic dynamics^25^. However, it is generally considered challenging to definitively demonstrate the presence of ecological chaos. Nevertheless, chaotic population dynamics have been convincingly found in a handful of well-controlled laboratory^26–29^ and field^30–32^ systems. Additionally, ecological chaos has recently been suggested to be an underappreciated and ubiquitous driver of abundance fluctuations across diverse populations^33^.

Despite the strength and near-universality of such large population abundance fluctuations, they are not believed to strongly affect the evolutionary dynamics of populations. Evolution is primarily driven by the dynamics of the relative frequency of alleles; if the population is experiencing strong abundance fluctuations, the allele frequencies will be unaffected if all alleles have synchronous abundance fluctuations. The primary source of stochasticity in allele frequency trajectories is generally thought to be genetic drift^16,35–37^. Classical genetic drift is a form of demographic stochasticity that arises from stochastic offspring number fluctuations, as represented in models such as the Wright-Fisher model^38^. Genetic drift is expected to have a relatively small impact on abundance fluctuations, especially at large population sizes. However, the prevalence of giant abundance fluctuations in ecological dynamics warrants further investigation into their potential evolutionary implications. We hypothesize that these giant fluctuations could drive large frequency changes in subpopulations with asynchronous, stochastic abundance fluctuations. This hypothesis challenges traditional evolutionary models by suggesting that large abundance fluctuations can sometimes influence relative genotype frequencies.

To investigate this, we turned to using genotypes isolated from the *E. coli* Long Term Evolution Experiment (LTEE). The LTEE is a well-known model system in experimental evolution, where several replicate *E. coli* populations have been propagated for over 70,000 generations, evolving in a simple daily dilution environment^39^. The daily dilution environment leads to repeated population bottlenecks, where only one out of every one hundred cells is propagated into the next day’s flask. This bottlenecking is expected to lead to result in genetic drift analogous to that described by the Wright-Fisher model. As a model system, we used two LTEE-derived strains that have coevolved with each other, referred to as *S* and *L*^40–43^. *S* and *L* diverged from each other early in the LTEE evolution, around 6.5k generations, where *S* emerged as an ecologically-distinct, but closely related genotype that partially invaded the initially *L*-dominated population^44,45^.

Ascensao et al. (2023)^34^ previously created random barcoded transposon knockout libraries of *S* and *L*, allowing them to track the frequency dynamics of many subclones of each strain within populations via amplicon sequencing. When they co-cultured the *S* and *L* libraries together, they saw that the total frequency of *S* relative to *L* fluctuated strongly (Figure1A). In contrast, the fluctuations of neutrally-barcoded variants of *S*, relative to the total *S* population were significantly more muted (Figure1B). The same observation holds true for *L* (FigureS1). Quantifying the strength of the observed frequency fluctuations, we see that the fluctuations between *S* and *L* are many orders of magnitude larger than fluctuations within-*S* (Figure1C). Additionally, the within-*S* fluctuations are similar to the variance expected from bottlenecking, thought to be the primary source of genetic drift in serial transfer environments.

**Figure 1.**
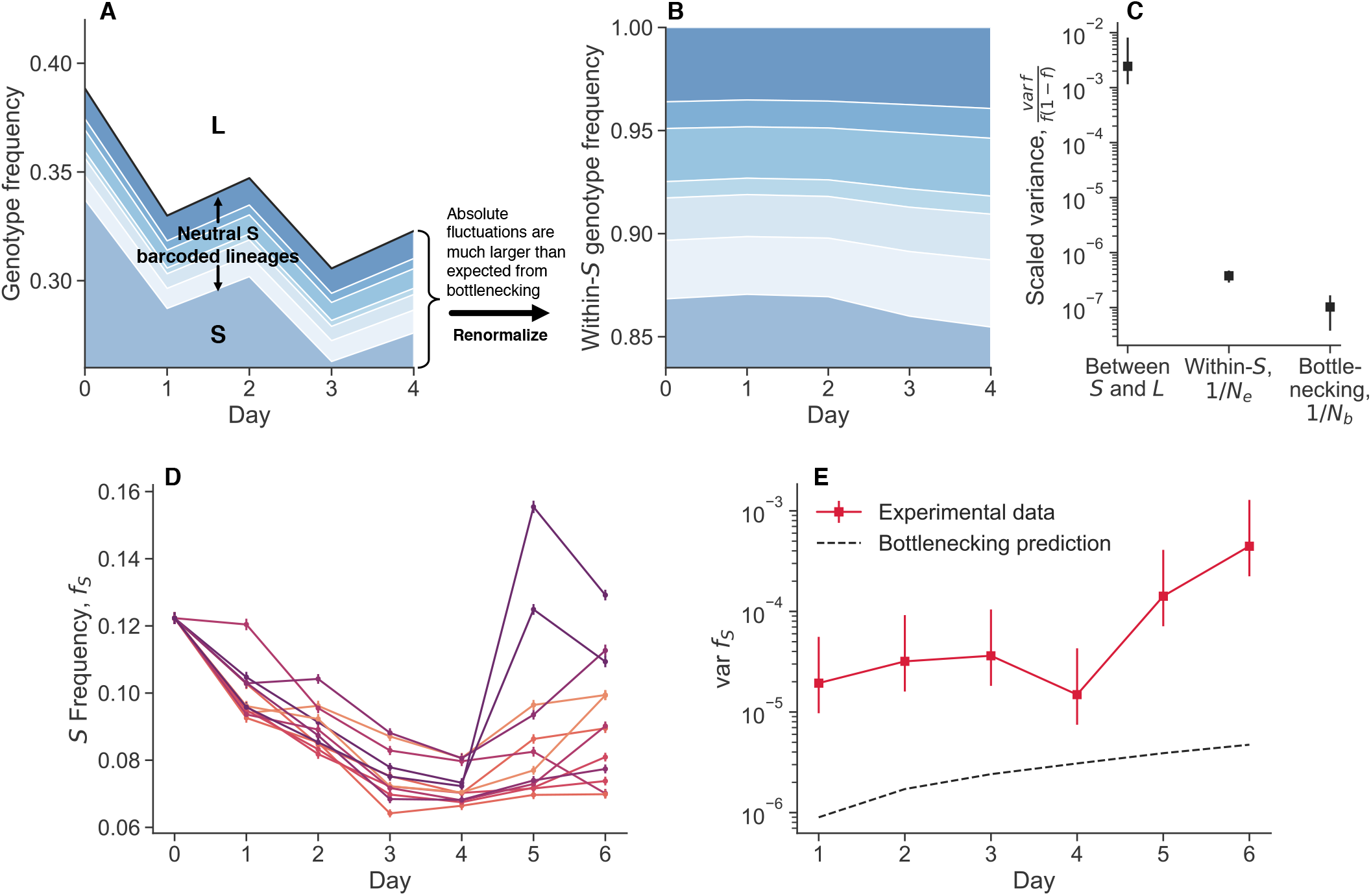
Observation of large genotype frequency fluctuations. (**A, B**) Barcoded libraries of *E. coli* strains *S* and *L* were propagated together in their native serial dilution environment (previously reported data^34^). In a Muller plot representation of lineage sizes, we see that the total frequency of *S* relative to *L* shows large fluctuations. However, neutral barcoded lineages within *S* show substantially smaller fluctuations relative to each other. (**C**) By quantifying the strength of fluctuations, we see that total frequency fluctuations between *S* and *L* are several orders of magnitude larger than fluctuations between neutral lineages and expected fluctuations from bottlenecking. (**D, E**) We propagated replicate cocultures of *S* and *L* strains together, after splitting them from the same mother culture at day zero. Even after one day of propagation, there is already more variance between replicates than expected from bottlenecking, and the variance accumulates over time. Note that the experiments in panels **A-C** and in **D-E** were performed at different culture volumes (800mL versus 1mL), but under the same daily dilution rate. All error bars represent 95% CIs.

We performed another coculture experiment with *S* and *L*, and measured relative frequencies with flow cytometry, a measurement technique orthogonal to amplicon sequencing. We propagated an *S*/*L* coculture, and then split the coculture into eight replicate cultures at day zero (Figure1D). We continued to propagate the replicate cultures separately, but in the same environment. After a single growth cycle, there is already more variance between replicates than would be expected from classical genetic drift, and it accumulates over time (Figure1E). Measurement noise cannot explain the magnitude of the variance, nor the fact that it tends to accumulate over time. We did not find these large fluctuations when we tested another, related pair of diverged genotypes in coculture, REL606 (the LTEE ancestor), and a strongly beneficial mutant, REL606 Δ*pykF* (FigureS2). Instead, we found that the variance accumulation was consistent with classical genetic drift. This indicates that not all non-neutral genotype pairs exhibit these giant fluctuations, and serves to provide additional evidence that there are no additional, unexplained sources of technical variance.

What is the source of these large observed fluctuations, and how do they behave? We find that these giant frequency fluctuations act differently compared to classical genetic drift, leaving a distinctive footprint on the population dynamics. We constructed an effective model that demonstrates how large random abundance fluctuations can arise from correlated offspring numbers between individuals. Giant frequency fluctuations originate when the offspring numbers of individuals *within* genotypes are more correlated than those *between* genotypes. We thus refer to such fluctuations as “decoupling noise”. Our analysis further uncovers that these offspring number correlations are primarily driven by underlying chaotic dynamics. These dynamics induce a fluctuating selection-like effect, which significantly influences the population’s evolutionary trajectory. Our findings indicate that decoupling noise is likely common in various biological populations, fundamentally impacting evolutionary and ecological dynamics. This study not only provides a deeper understanding of the mechanisms behind population fluctuations, but also underscores the importance of updating traditional evolutionary theories to integrate these dynamic complexities.

## Results

### Empirical fluctuation scaling measurements

We first aimed to determine if the large fluctuations we have observed behave in the same way as classical genetic drift. Under neutral drift, the variance in genotype frequency after one generation, var *f*, will depend on the mean frequency, var *f* = ⟨*f* ⟩ (1 − ⟨*f* ⟩) */N*_*e*_^38^. At low frequencies, *f* ≪ 1, the variance will scale linearly with mean, var *f* ≈⟨ *f* ⟩ */N*_*e*_. Here, the brackets ⟨·⟩ represent the mean of a random variable. Deviations from the predicted scaling behavior would indicate fluctuations that do not arise from classical genetic drift.

We sought to measure the mean-variance scaling relationships of population fluctuations in the *S* and *L* coculture system, measuring population abundances and genotype frequency via flow cytometry (Figure2A). Briefly, we initially grow each genotype in monoculture for several serial dilution cycles, before mixing the two genotypes together at a defined frequency. After one more growth cycle, we split the “mother culture” into sixteen replicate daughter cultures, all grown in the same environment. After another growth cycle, we take flow cytometry measurements of all daughter cultures, and then compute the mean and variance across the biological replicates derived from the same mother culture. We varied the initial frequency of the minor genotype, *S*, over about two orders of magnitude.

**Figure 2.**
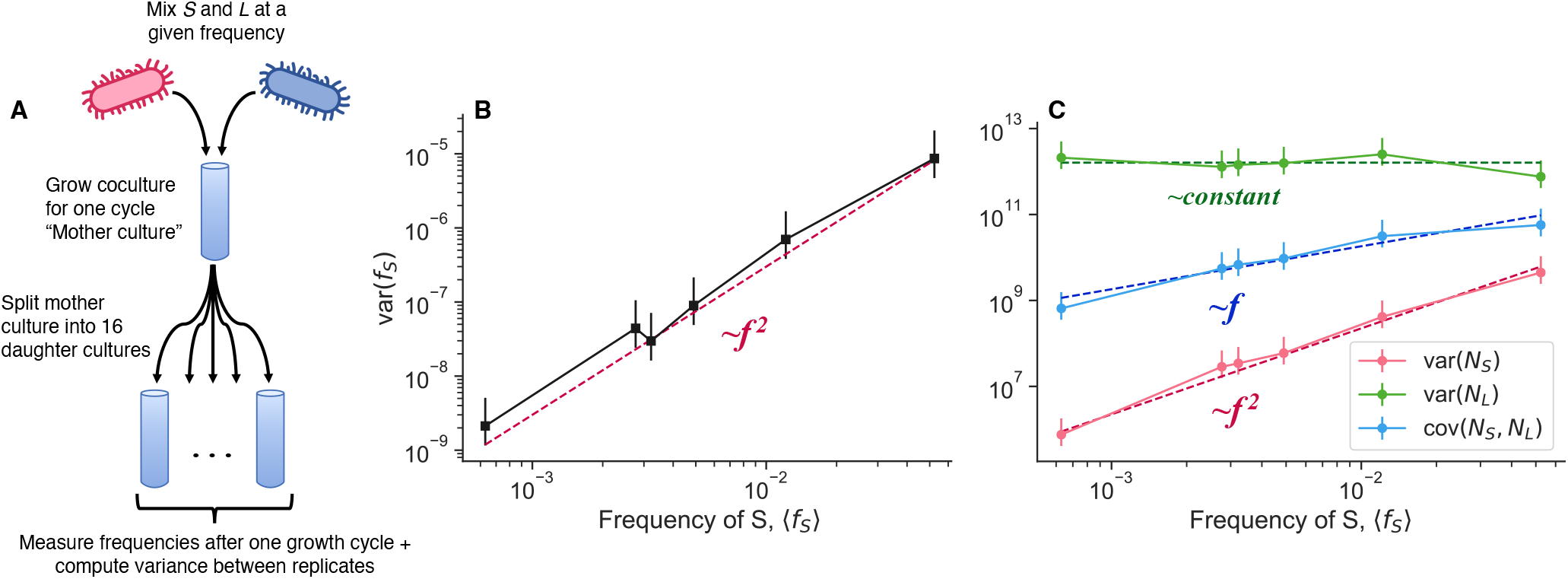
Empirical scaling of genotype frequency and abundance fluctuations. (**A**) After cocultering *S* and *L* together at defined relative frequencies, we split the mother cultures into 16 replicate daughter cultures. We then grew the cultures for another cycle, and measured the mean and variance across cocultures. We measured (**B**) variance of genotype frequency along with (**C**) variance and covariance of absolute abundance, both as a function of mean *S* frequency. The points represent experimental measurements, and the dashed lines are fitted lines with the indicated scaling. Error bars represent 95% CIs.

Once we calculate the variance across replicate daughter cultures, we see that the variance of *S* frequency scales approximately like var 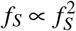 (Figure 2B). We also measured the scaling behavior of the variance of the absolute abundance of *S* and *L*, var(*N*_*S*_) and var(*N*_*L*_) respectively, and the covariance between the two genotypes cov(*N*_*S*_, *N*_*L*_) (Figure 2C). Importantly, as with frequency, we measured the total abundance at the end of the growth cycle; we were not considering intra-cycle changes in abundance. The abundance of *S* scales like var 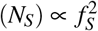, *L* abundance stays approximately constant, and the covariance scales linearly, cov(*N*_*S*_, *N*_*L*_) ∝ *f*_*S*_. These observations also deviate from the prediction of classical genetic drift–the variance of *S* abundance should scale linearly, var(*N*_*S*_) ∝ *f*_*S*_, and the covariance should be zero or negative (if the population has a set carrying capacity), cov(*N*_*S*_, *N*_*L*_) ≤ 0. Furthermore, the data indicate that it is not the case that the fluctuations predominantly arise from only one genotype–*S* and *L* abundance fluctuations are of about the same magnitude, with *S* potentially fluctuating slightly more by a factor of order one (Figure S4). Additionally, we measured total abundance fluctuations as a function of initial population sizes, by varying the volume of the culture while holding the dilution rate constant (Figure E8). We also found that var(*N*) ∝ *N*^2^, in both monoculture and coculture conditions. This indicates that the power law-scaling abundance fluctuations are present even in the absence of coculture conditions.

Together, these data clearly indicate that the large frequency fluctuations we see cannot be explained by classical genetic drift. Now a new question arises: what type of process is generating the observed fluctuation scaling behaviors?

### Effective model of population fluctuations

We would like to understand how genotype frequency and abundance fluctuations may arise from variation in individuals’ offspring number (SI section S2.1). We first consider a simple population consisting of two genotypes, where all individuals are identical, except for a neutral (genetic) difference to distinguish between the types. There are initially *N*_*A*_ and *N*_*B*_ individuals of genotype A and B, respectively. Each individual *i* gives rise to 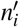 net offspring in the next generation, where var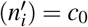. We define the total population abundance as *N*_*tot*_ = *N*_*A*_ + *N*_*B*_, and frequency of genotype A as *f* = *N*_*A*_*/N*_*tot*_. We can then show that the variance of total population abundance and frequency in the next generation will be,

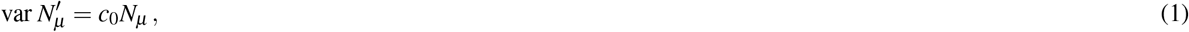

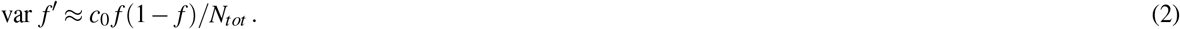

Thus, we see that we recover the variance relationships for fluctuations due to classical genetic drift, and we explicitly see that classical genetic drift arises from the sum of independent offspring number fluctuations. How can we now extend or generalize this model? One simple extension would be to allow individuals within a population to have correlated offspring numbers. This could arise due to a randomly fluctuating environment which induces transient opportunities (or perils) for reproduction, causing offspring numbers to be coordinated across individuals. We can introduce a new covariance parameter, cov 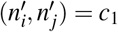, for different individuals, *i* ≠ *j*. The variance in population abundance and frequency becomes,

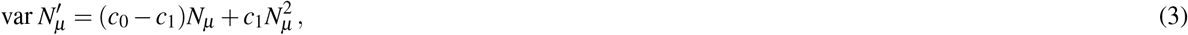

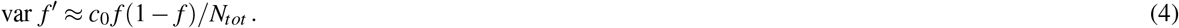

The form of this total population abundance variance scaling has been previously noted^46,47^. The abundance variance now scales linearly at low abundance, but shows quadratic growth at higher abundance. Power law mean-variance scaling of population abundance has been widely observed in ecology, where it is known as Taylor’s power law^1–5^. The variance in frequency stays the same as the case with uncorrelated offspring numbers; intuitively, this is because even though the abundance can strongly fluctuate due to correlated offspring numbers, the population sizes of the two genotypes will fluctuate in sync, so there will be no net effect on frequency fluctuations. However, this will only be the case if the offspring number fluctuations are correlated in precisely the same way with individuals of the same genotype and with individuals of a different genotype. Thus, we propose a yet more general model where the two genotypes are not necessarily identical, and the covariance parameters can depend on the genotypes considered (Figure 3). We consider the offspring number covariance between two individuals, where 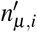 represents in the number of net offspring from individual *i*, which belongs to genotype *µ*,

**Figure 3.**
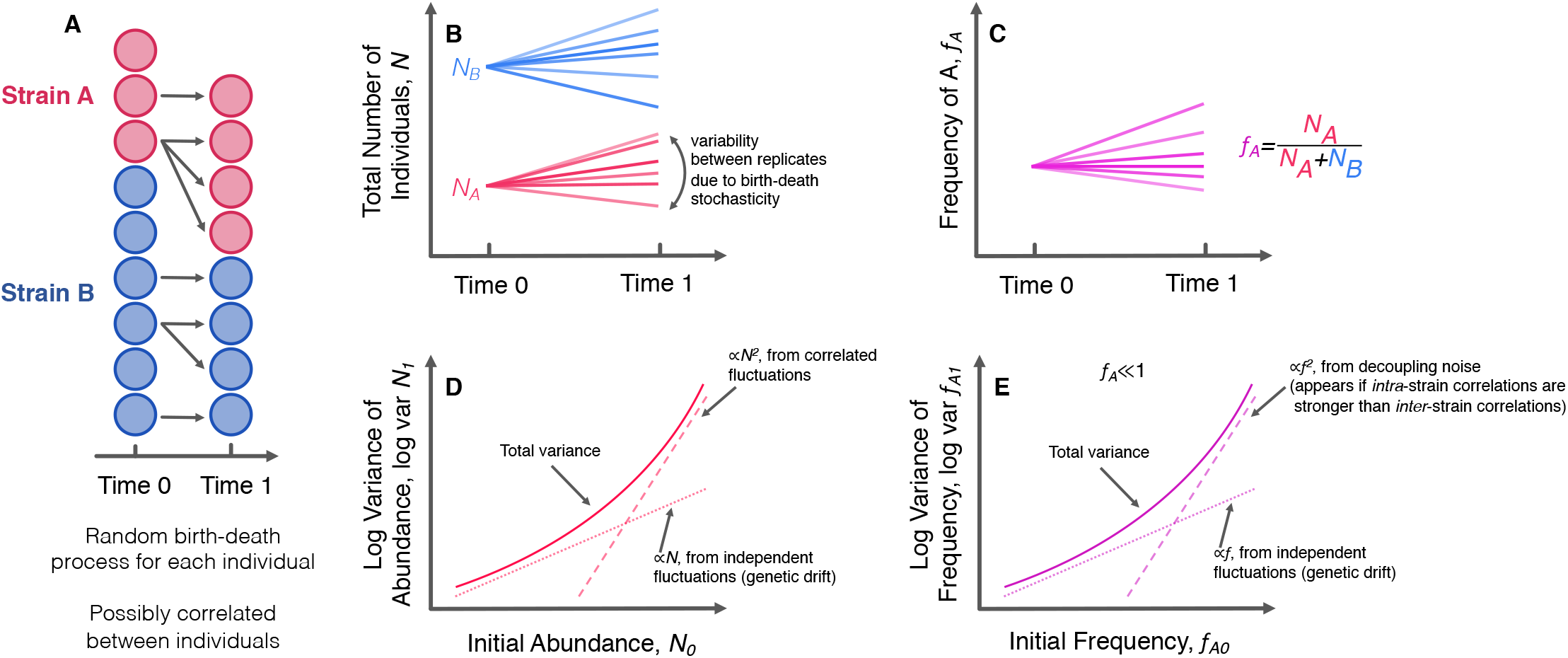
Model of population fluctuations. (**A**) We consider a population of individuals, subdivided into two genotypes. At time zero, there are a specified number of individuals in each genotype. After some period of time, each individual has left behind some random number of descendants, drawn from a distribution that may be correlated between individuals. (**B, C**) The abundance of populations and relative frequencies of genotypes will generally differ across different replicate populations split from the same mother culture, due to the random nature of the process. (**D, E**) Our model suggests specific scaling behaviors for both the variance of a total number of individuals in each genotype and variance of the relative frequencies. Specifically, there is a linearly scaling component caused by independent fluctuations (classical genetic drift), and a quadratically scaling component caused by fluctuations correlated between individuals. The quadratically scaling fluctuations will appear in abundance trajectories if there are correlated fluctuations, but they will only appear in frequency trajectories if *intra*-genotype correlations are stronger than *inter*-genotype correlations.

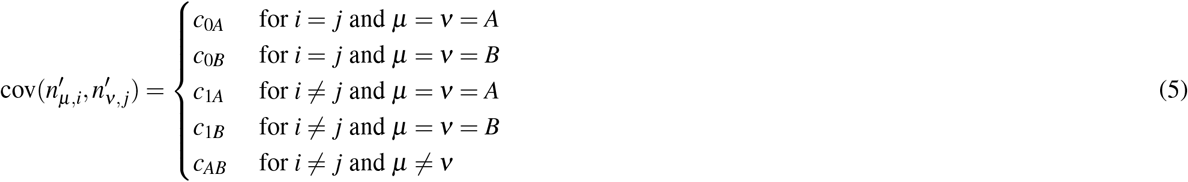

The abundance variance does not change (equation 3), and the covariance between the abundance of the two genotypes along with the frequency variance becomes,

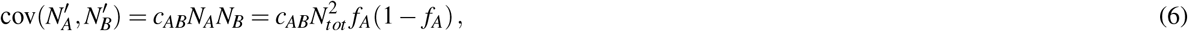

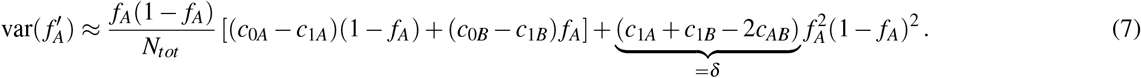

The new composite parameter *δ* quantifies the degree of decoupling between the two genotypes. If the genotypes are identical such that *c*_1*A*_ = *c*_1*B*_ = *c*_*AB*_, then the quadratically-scaling fluctuations will vanish. These fluctuations will only appear if the offspring numbers of individuals *within* a genotype are more correlated with each other compared to individuals *between* genotypes. Similar forms for the frequency variance were found by Takahata et al. (1975)^48^ and Melbinger and Vergassola (2015)^21^; however, we consider the more general formulation where all five covariance parameters may differ from each other, and our model can be derived in a more generic way. Our model is readily extensible–we can expand our results to the more general case of a population consisting of *m* different genotypes (SI section S2.3).

We note that if we consider the case where *c*_0*A*_ − *c*_1*A*_ = *c*_0*B*_ − *c*_1*B*_, then we can simplify equation 7,

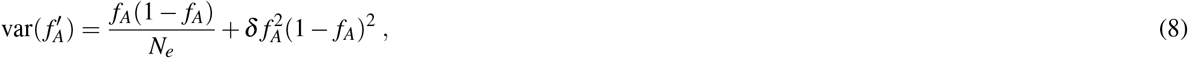

where the effective population size is defined as *N*_*e*_ = *N*_*tot*_*/*(*c*_0*A*_ − *c*_1*A*_). This form of the variance of genotype frequencies clearly shows that it is composed of two components. The first component arises from independent fluctuations of individuals, linearly scales with frequency, and corresponds to classical genetic drift. The second, quadratically scaling part arises when offspring number fluctuations between genotypes are decoupled to a degree, thus we refer to it as “decoupling noise”.

Our model is an *effective* theory that purports to describe mean-variance scaling behaviors through covarying offspring numbers; such covariances may arise from a number of different underlying dynamical processes. Our model shows that abundance fluctuations with a power-law exponent of two *imply the existence of correlations between the reproductive success of individuals*. Said another way, if a given dynamical system displays such abundance fluctuation scaling, the source of the fluctuations can be interpreted as offspring number correlations. Analogously, genotype frequency fluctuations with a power-law exponent of two imply that genotypes have decoupled offspring number correlations. As previously mentioned, genotype frequency fluctuations with a power-law exponent of two can arise from fluctuating environmental conditions, which has been referred to as “fluctuating selection”^21,48^. However, fluctuating selection is not the only mechanism that can cause decoupling noise to appear. Various additional mechanisms^2–5^ can cause the correlated abundance fluctuations that scale like var *N* ∝ *N*^2^, including inherently chaotic dynamics^23,24^ and spatial effects, such as aggregation and dispersal^1,22^.

The variance and covariance scaling behaviors in equations3,6,7are all consistent with the experimentally measured scaling relationships (Figure2B-C). It appears that the measured range lies in the regime where the linearly-scaling component (classical genetic drift) is negligible compared to the effect of the correlated offspring number fluctuations. There is some evidence that the lowest data point in Figure2B may fall into the cross-over between the linear and quadratic regimes, but it is not completely clear (FigureS6). The correlation between *S* and *L* abundance fluctuations at high frequencies 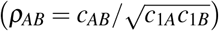 is over 90%, demonstrating that a slight decoupling in correlated fluctuations between genotypes is sufficient to generate noticeable decoupling noise. Overall, the quantitative agreement between the experimental data and our model points to the presence of correlated offspring number distributions in the *S*/*L* system. However, the origin of such correlated offspring number fluctuations is still not clear.

### Within-cycle growth measurements reveal chaotic dynamics

Populations derived from the *E. coli* LTEE are grown in a serial dilution, glucose minimal media environment, such that the populations are transferred at a 1:100 dilution into fresh media every twenty-four hours. This set-up creates a seasonally-varying environment, where the populations are switching strategies throughout a cycle as it proceeds from feast to famine and back again^42,49,50^. We reasoned that the within-cycle dynamics of replicate cultures could help to reveal the origin of decoupling noise. We find evidence that underlying chaotic dynamics are the source of the offspring number correlations between individuals.

We measured the population dynamics of *S* and *L* coculture over the course of the twenty-four hour growth cycle. In a protocol similar to the one used in first Results section, after several initial monoculture growth cycles, we mixed *S* and *L* such that *S* initially occupies around 6% of the population. After one more growth cycle in coculture, we split the mother culture into multiple, independent replicate daughter cultures, and started to take population measurements over defined time increments via flow cytometry. We grew all of the daughter cultures together in a shaking 37^°^C water bath, to minimize the effects of any possible environmental fluctuations. We measured the dynamics of the first eight hours and those of the last sixteen hours separately (on different days), because we found that the two periods had distinct experimental design requirements. The first eight hours (i.e. during exponential phase) is the period of the fastest dynamics, so we had to use both fewer biological replicates and a more dense sampling strategy. The last sixteen hours corresponds to stationary phase, where the dynamics are relatively slow.

We first look at the within-cycle dynamics of *S* frequency, *f*_*S*_ (Figure4A). We see relatively complex, out of steady-state dynamics, especially in the first eight hours. As previously described^51^, the dynamics can be explained by differences in lag time, exponential growth, and stationary phase behavior. The frequency of *S* initially increases because it “wakes up” from lag phase earlier than *L*. However, *L* has a higher growth rate on glucose, so *f*_*S*_ starts to decline once *L* wakes up. Then after a transition period, *f*_*S*_ starts to increase again due to a stationary phase advantage and better growth on acetate^42,51,52^.

We quantify the variance between replicates, and observe that there are periods of increased variance in approximately the first 5 hours, and the last 7 hours (Figure4B). The increased variance between replicates in the first 5 hours may be caused by the fast dynamics of exponential phase, but its origin is not definitively clear. The fact that the variance drops close to zero by eight hours, instead of accumulating, suggests a non-biological origin to explain the initial variance. In contrast, after a period where the variance does not increase much, we see a steady accumulation of variance later in the timecourse. Specifically, the variance appears to be increasing exponentially at a constant rate, from around 7 hours to the end of the 24 hour cycle (Figure 4C). This pattern is not consistent with technical sources of noise. The data fits an exponential trajectory better than linear or various other non-linear models (FigureS7). Biological replicates continually fluctuate and change their frequency rank order until the end of the time course (FigureS9)–their relative position is not “frozen in” early in the time course.

**Figure 4.**
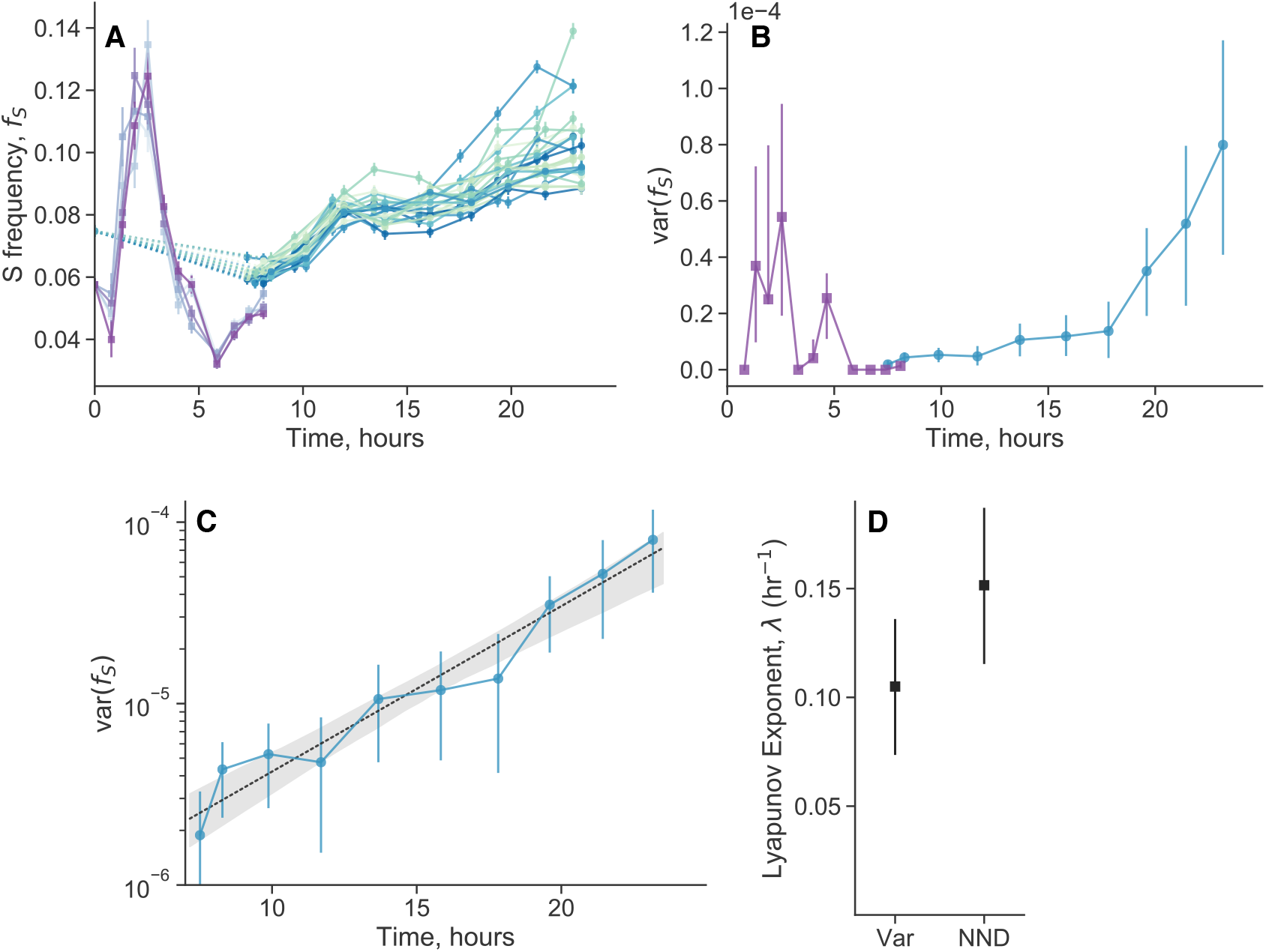
Within-cycle chaotic dynamics of genotype frequencies. (**A**) After splitting cocultures of *S* and *L* into multiple biological replicates, we measured genotype frequencies over the course of a twenty-four hour cycle (purple lines: 5 replicates; blue/green lines: 23 replicates). Each line represents a biological replicate. (**B**) Quantifying the variance across replicates over time, we see that the variance peaks both in the first 5 hours, and at the end of the cycle. However, the initial variance is not maintained beyond the first 5 hours, suggesting that the later accumulation of variance is the primary contributor to the decoupling noise. (**C**) We plotted the variance (after 7 hours) on a semilog scale, revealing that the variance appears to increase exponentially over time. The black dashed line represents the exponential fit. Exponentially increasing variance between initially close replicates is indicative of chaotic dynamics. (**D**) Lyapunov exponents calculated from the trajectories after 7 hours, inferred from the exponential fit from the variance (Var) and from the nearest-neighbor distance (NND) method. All error bars represents 95% CIs.

Exponentially increasing variance between replicates that are initially close to each other is indicative of chaotic dynamics. Chaotic dynamics are classically indicated by extreme sensitivity to tiny perturbations (and a bounded phase space), such that small differences in the initial conditions exponentially increase over time. The observation of exponentially increasing variance is equivalent to the observation of pairwise exponential divergence (SI sectionS4.1).

We used another standard method to detect chaotic dynamics and infer the largest Lyapunov exponent, based on changes in nearest-neighbor distance (NND)^53^. We inferred a significantly positive Lyapunov exponent (*p* = 0.006), which is consistent with the Lyapunov exponent estimated from the exponentially increasing variance (Figure 4D). The inverse of the Lyapunov exponent (“Lyapunov time”) represents a characteristic timescale of the system, effectively representing how long a system will appear to be predictable. The Lyapunov time is approximately 5-10 hours, implying that trajectories would appear to be stochastic on longer timescales. Thus, the overnight timecourses were necessary to reveal the chaotic dynamics, because the system shows a relatively fast Lyapunov time of approximately 5-10 hours, so timepoints taken every 24 hours appear effectively stochastic.

Together, these data suggest that decoupling noise originates from underlying chaotic dynamics. Chaotic dynamics cause individuals in a population to coordinate their birth/death rates (in an unpredictable way), implying the presence of effective offspring number correlations, and thus leading to mean-variance power law exponents of two. More generally, Ballantyne (2005)^24^ argued that Taylor’s power law with exponent of two will occur in any deterministic population dynamics model under two conditions: (i) the population size continues to fluctuate over time and (ii) the population growth rate remains constant. The Taylor’s power-law scaling is a simple consequence of the ability to rescale such deterministic systems by a constant factor *k*, which implies that the coefficient of variation remains constant,

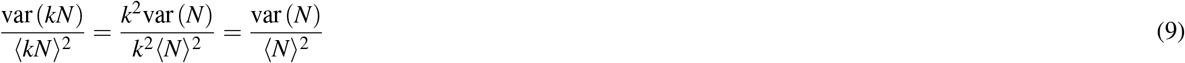

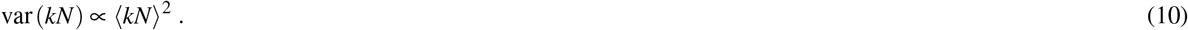

Decoupling noise can then arise when the chaotic fluctuations between two populations are (to some degree) asynchronous.

To further elucidate how chaotic dynamics can give rise to effective correlated offspring numbers, and decoupling noise, we simulated the chaotic dynamics in a simple two-genotype model (Figure E9). The model we analyze here is not intended to represent the underlying dynamics of the *S*/*L* system, but rather serves to demonstrate generic properties of chaotic dynamics. In alignment with our prior expectations, we see that Taylor’s power law holds, with var *N* ∝ *N*^2^. Additionally, decoupling noise appears across a range of different coupling strengths between the two genotypes, var *f* ∝ *f* ^2^.

### Extrinsic versus intrinsic decoupling noise

Most prior experiments that we performed focused on dynamics across one 24 hour growth cycle. However, in both evolution experiments and natural populations, evolutionary and ecological dynamics occurs across many growth cycles or much longer time periods. In prior experiments that we performed over a single growth cycle, all replicates shared the same mother culture and experienced the same environment, controlling for the effects of (potentially subtle) environmental noise and between-day memory-like effects. However, in principle, both environmental noise and memory-like effects could impact the overall effective strength of decoupling noise (SI section S2.2). Both effects would induce offspring number correlations that would also cause correlations between replicate cultures grown in the same environment, and those that shared a mother culture. In an analogy to gene expression noise^54^, we refer to decoupling noise that is not correlated between replicate cultures as “intrinsic noise” (Figure 5A), which is the effect driven by inherently chaotic dynamics that we’ve been focusing on in previous sections. We use the term “extrinsic noise” for decoupling noise that is induced by shared environmental fluctuations (Figure 5B); this component has also been referred to as “fluctuating selection”^21,48^. We thus sought to estimate the total strength of decoupling noise in our system by isolating the effects of shared mother cultures, intrinsic noise, and extrinsic noise.

**Figure 5.**
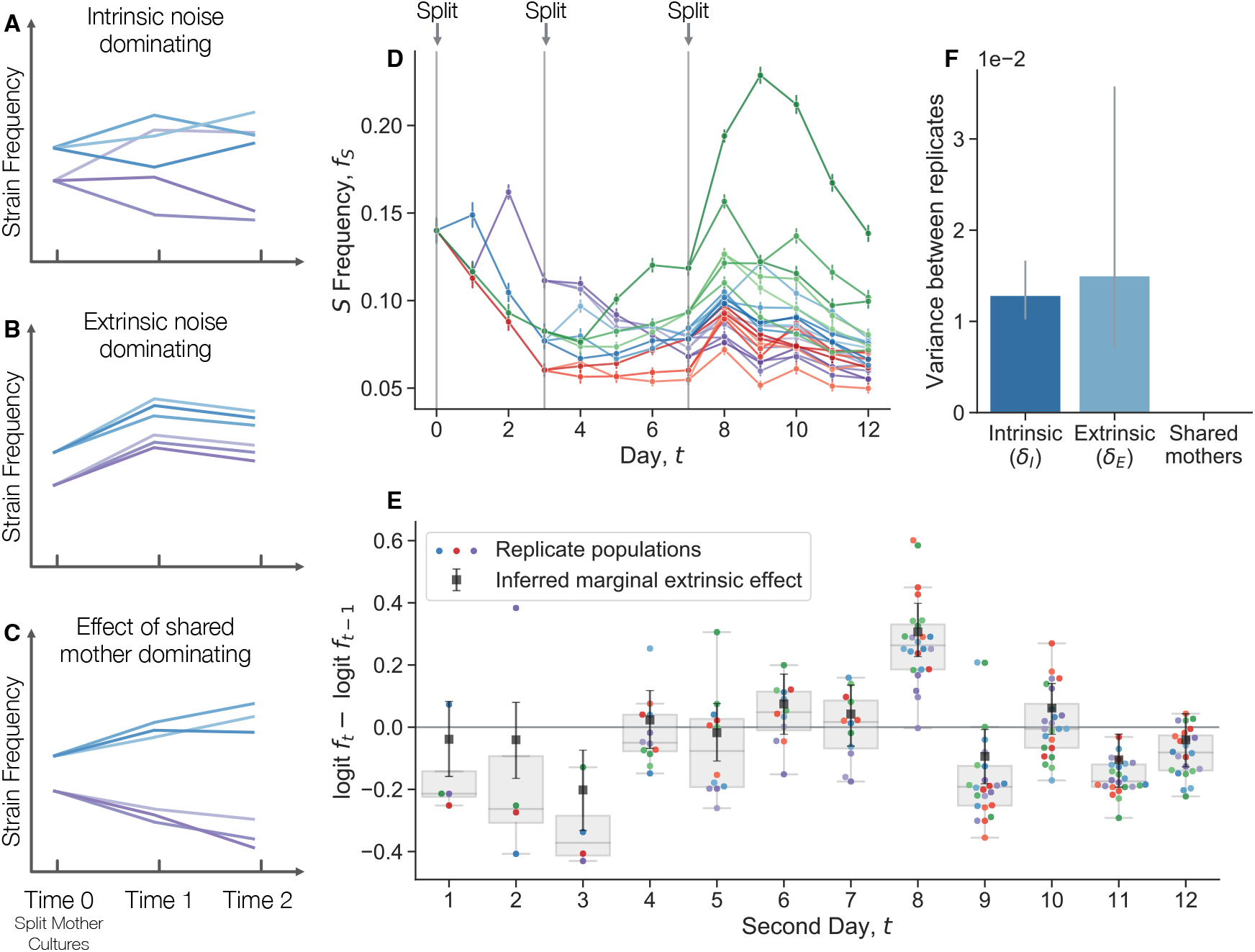
Quantifying the relative strength of intrinsic noise, extrinsic noise, and the effect of sharing mother cultures. (**A-C**) Different sources of decoupling noise leave different signatures in genotype frequency time courses, arising from different sources of offspring number correlations. Schematics show two initial mother cultures that are each split into three replicate daughter cultures at time zero. (**A**) Intrinsic decoupling noise causes frequency fluctuations that are uncorrelated between replicates. (**B**) Extrinsic noise causes correlations in replicate cultures sharing the same environment. (**C**) Memory-like effects would cause (temporary) correlations between cultures split from the same mother culture. (**D**) We performed an experiment where we cocultured *S* and *L* together, then split the coculture into four replicate cultures on day 0. We continued to propagate the cultures, and subsequently split each culture into more replicate cultures on days 3 and 7. (**E**) We computed the change in logit frequency from one day to another (relative, extrinsic fitness effect) for each replicate population. Colors for each population are consistent with panel **D**. The black squares represent the inferred extrinsic “fitness effect” for each day pair, controlling for the effect of frequency-dependent selection, shared mothers, intrinsic noise, and measurement noise. (**F**) We developed a model to partition variance of frequency noise, reporting the posterior median. All error bars represent 95% CIs.

We conducted an experiment where we propagated *S*/*L* cocultures over several days in the same environment, serially splitting cultures into new replicates at three different timepoints (Figure 5D). We see that the frequency of *S* initially declines across replicate populations, owing to its frequency dependent fitness effect (Figure S10). The populations stabilize around their equilibrium frequency, but continue to fluctuate. We quantified the change in logit frequency from one day to the next for each replicate population (Figure 5E), i.e. the between-day “fitness effect”. We see that there are several days where the change in frequency is noticeably correlated across replicates; for example, from days 7 to 8, most replicates appear to increase in frequency, even though they were at a large range of different frequencies. This seems to indicate that there are significant extrinsic fitness fluctuations, putatively caused by subtle environmental noise. However, there are various possible contributions to fluctuations at each day, including measurement noise, extrinsic and intrinsic decoupling noise, frequency-dependent fitness, and any memory-like effects from sharing a mother culture. To pull apart the contributions of each effect, we built a Bayesian hierarchical model and fit it to the data (see Methods section “Extrinsic fluctuations and splitting cultures”). Briefly, we model the change in logit-transformed frequencies as Gaussian random variables, that can be affected by the aforementioned contributions. We see that intrinsic and extrinsic noise contribute roughly the same level of variance to frequency fluctuations (Figure 5F). In contrast, there is little detectable effect of sharing a mother culture (upper bound of the 99% credible interval is ∼ 10^−15^), indicating that any memory-like effects are overshadowed by inherent and environmental fluctuations. The large extrinsic fluctuations are perhaps surprising, given that the cultures were maintained in the same temperature and humidity-controlled incubator for the duration of the experiment. Similar sensitivity to putatively subtle environmental fluctuations has been observed in other experiments^55^. This sensitivity is likely caused by the underlying chaotic dynamics of the process, which can exponentially amplify minor environmental differences.

Both intrinsic and extrinsic decoupling noise appears to be present in other, unrelated experimental systems. Venkataram et al. (2016)^56^ used a barcoding system to track frequency trajectories of many adaptive variants of *S. cerevisiae* yeast. They also find large frequency fluctuations when adaptive variants are at high frequencies. Venkataram et al. cultured populations together, and then split the cultures into three biological replicates at time point 1. We exclude batch 2 from our analysis because it only had two replicate cultures per time point, compared to three replicate cultures for batches 1, 3, and 4. We computed the frequency variance between biological replicates after one growth cycle apart (Figure 6A), leveraging the presence of many adaptive barcoded clones to average across the clones to obtain a more precise mean-variance relationship. We pooled all barcoded mutants together in this analysis, and thus we are averaging over the effects across genotypes. We see that the there is an uptick in the variance at high mean frequencies. Specifically, variance in frequency between biological replicates scales approximately like ∼ *f* at low frequencies and ∼ *f* ^2^ at high frequencies, again consistent with equation 7. An unconstrained power-law fit on the raw data yields a power-law exponent of 1.14 ±0.05 at low frequencies, and 2.0 ± 0.08 at high frequencies. We also computed the relationship between the mean variance and the variance of log frequencies (Figure 6A inset), which is also consistent with our model, as we would expect a constant mean-variance relationship at high frequencies (the log transformation acts as a variance-stabilizing transform). To investigate if the putative decoupling noise accumulates over time, we estimated the mean-squared displacement (MSD) of the log-frequency trajectories (of barcodes with high mean frequency), correcting for measurement noise and fitness effects (Figure 6B). If decoupling noise is indeed causing the observed quadratic variance scaling at high mean frequencies, the MSD should increase at an approximately linear rate, with the slope corresponding to *δ*_*I*_. Consistently with our prediction, we see that the MSD increases with increasing time increments across all batches. The strength of intrinsic decoupling noise is approximately the same across the two major classes of adaptive mutants, adaptive haploids and diploids (Figure E10); virtually all high-frequency barcodes are adaptive mutants. If two clones were identical, we would expect that decoupling fluctuations would affect them in the same way, and thus their frequency displacements would be perfectly correlated. Indeed, we see that if two clones are of the same mutant class, their fluctuations are more correlated on average than between clones of different mutant classes (Figure E11).

**Figure 6.**
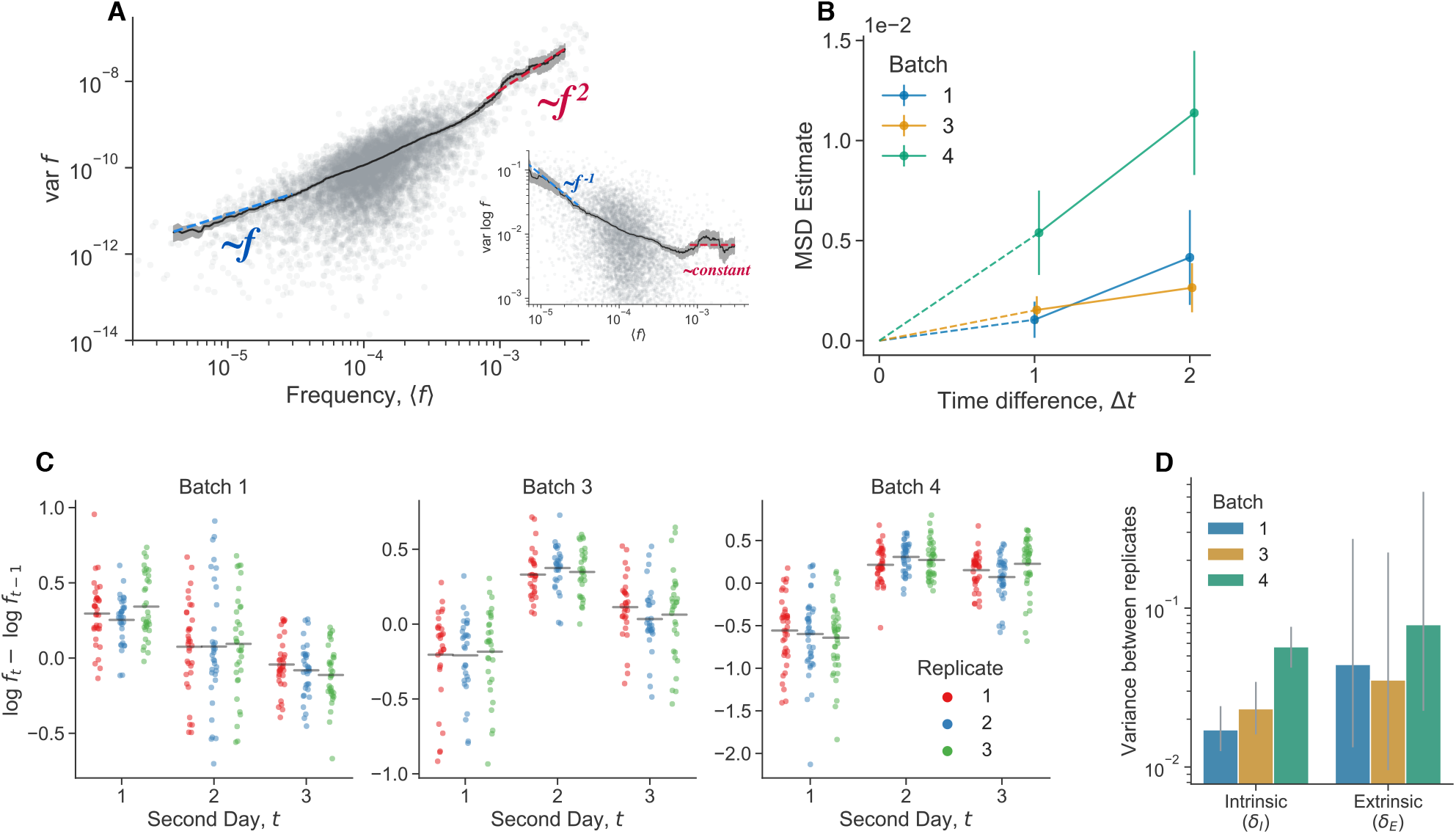
Extrinsic and intrinsic decoupling noise found in barcoded *S. cerevisiae* populations. Data reanalyzed from Venkataram et al. (2016)^56^. (**A**) We computed the barcode frequency variance between three biological replicates, one growth cycle after the culture was split into the replicates. We included all barcoded mutants in the analysis. Inset shows a different representation of the same data, computing the variance of the log-frequencies. The grey points represent the variance of individal barcoded mutants across the three biological replicates. The black line represents a rolling average. Colored dashed lines represent the indicated scaling. (**B**) Estimate of the mean squared displacement (MSD) of the log-frequencies (restricted to barcodes with high mean frequency). (**C**) Change in log-frequency from one day to another (relative, extrinsic fitness effect) for each barcode in each batch (again, restricted to high frequency barcodes). The gray line represents the mean displacement for each replicate at each timepoint. (**D**) Estimated contribution of both intrinsic and extrinsic decoupling noise to frequency fluctuations. All error bars represent 95% CIs.

Similarly to the previously presented data (Figure 5E), we investigated the effect of extrinsic noise by plotting the log displacement of high-frequency barcodes over time (Figures 6C). We see that the mean displacement of barcodes is often correlated at time points, in a way that is consistent within batches, but not between batches. This is potentially a signal of extrinsic decoupling noise. To quantify the relative strength of intrinsic versus extrinsic decoupling noise, we employ a similar Bayesian hierarchical model to the one previously presented (Figures 6D). We again infer relatively large strengths of both intrinsic and extrinsic decoupling noise. The inferred strength of extrinsic decoupling noise has wide, uncertain posterior across all batches, which is due to the fact that there are a small number of timepoints per batch. However, the error bars provide a lower bound on plausible values of *δ*_*E*_, which is of the same magnitude as the inferred strength of intrinsic noise. Together, these data provide evidence for the presence of both intrinsic and extrinsic decoupling noise in an experimental barcoded yeast system.

The effect of extrinsic fluctuations can easily be incorporated into our model (SI section S2.2). Specifically, if the environment has a negligible autocorrelation time, the total decoupling parameter is simply the sum of the intrinsic and extrinsic components, *δ* = *δ*_*I*_ + *δ*_*E*_. Significant environmental autocorrelation times could lead to more complex dynamics^57^. In principle, the extrinsic component can be altered easily by changing the rate/amplitude of environment fluctuations, while the intrinsic component depends on the inherent (chaotic) dynamics of the system.

### Evolutionary implications of decoupling noise

As fluctuations arising from classical genetic drift have strong effects on evolutionary outcomes, it is reasonable to expect that the presence of decoupling noise may also have evolutionary implications. To study the implications of decoupling noise, we consider our model in the diffusion limit, under constant selection (equation S20). Previous studies^21,48,58,59^ have investigated similar stochastic processes, but we consider the more general form where all five covariance parameters may differ from each other. We obtain analytical results for the fixation probability and the site frequency spectrum, and compare those results to simulations.

We consider a mutant with fitness effect *s* and decoupling parameter *δ* = *c*_1*A*_ + *c*_1*B*_ − 2*c*_*AB*_, defined in terms of the offspring number covariances. In this section, we focus on the case where the strength of genetic drift is the same between the two genotypes,

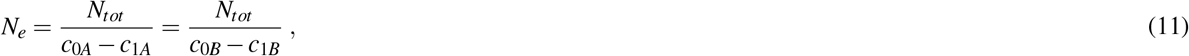

so that we can use one parameter, *N*_*e*_, to encapsulate the strength of genetic drift. However, we can generalize beyond this case (SI section S3).

### Fixation probability

We examine the fixation probability of a mutant in a two-genotype system (Figure 7A). We derive a general closed-form expression for the fixation probability as a function of the initial mutant frequency *f*_0_ (SI section S3.1). Notably, the fitness effect appears only within a new compound parameter, which we term the effective fitness effect, *s*_*e*_ = *s* +(*c*_1*B*_ − *c*_1*A*_)*/*2 (Figure S14).

**Figure 7.**
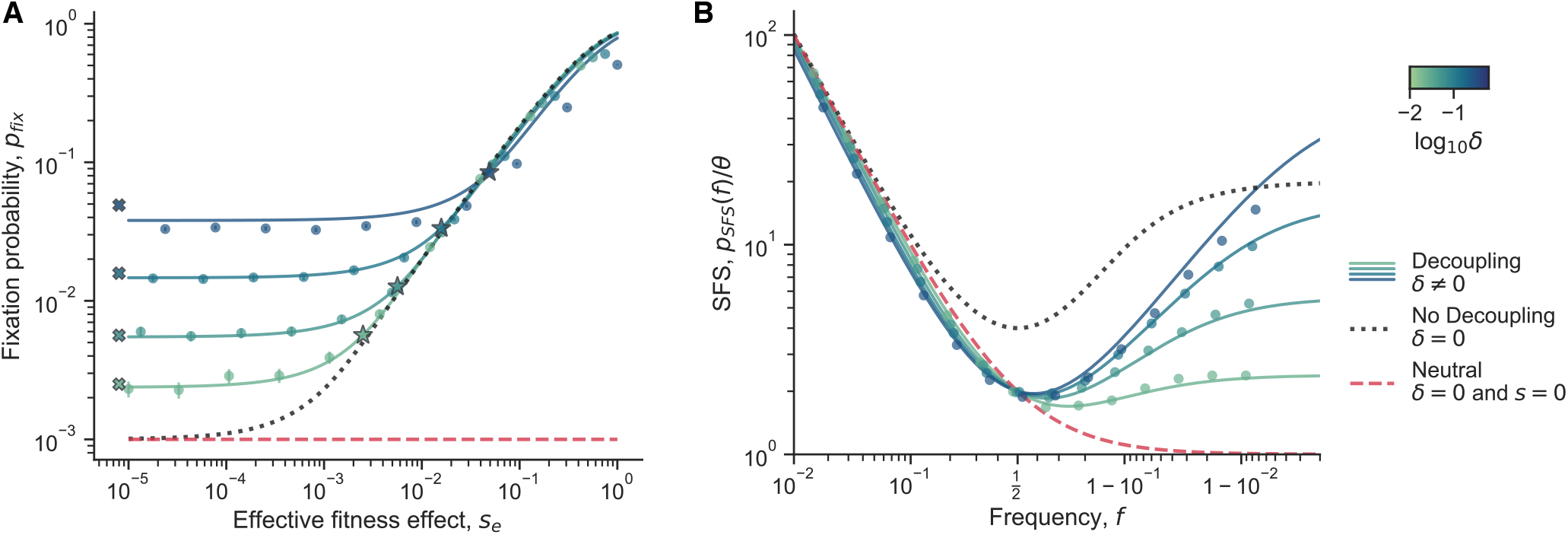
Theoretical evolutionary implications of decoupling noise. (**A**) The fixation probability of a beneficial mutant, as a function of its (effective) fitness effect, *s*_*e*_. Here we focus on the case that *c*_1*A*_ = *c*_1*B*_, so that *s*_*e*_ = *s*. The star markers show the location of 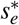 (equation 12), the approximate transition point between the two regimes of *p* _*fix*_. The *x* markers represent the asymptotic fixation probability at low *S* (equation 13). (**B**) The site frequency spectrum as a function of the mutant frequency (*logit* x-axis scale). In both plots, the blue-green solid lines represent full analytical solutions for dynamics with decoupling noise, across different values of *δ*. The round markers show simulation results; error bars represents 95% CIs. The black dotted lines represent the case where there are no decoupling noise, but there is still (constant) natural selection. The red dashed lines represents the case where there is neither decoupling noise nor natural selection. Across both plots, *c*_1*A*_ = *c*_1*B*_, *ρ*_*AB*_ = 0.5,*N*_*e*_ = 10^3^. In (**A**), *f*_0_ = 10^−3^. In (**B**), the solid lines show the case where *s* = 0 but *δ*≠0 and the black dotted line shows the case where *s* = 0.01 and *δ* = 0.

In the limit of weak decoupling noise, *δ* ≪ 1*/N*_*e*_, we find that *p* _*fix*_ reduces to the classical fixation probability of a mutant under constant selection^38^, as expected (Figure 7A; black dotted line).

We then examine the limit where (i) decoupling noise is much stronger than genetic drift, *δ* ≫ 1*/N*_*e*_, and (ii) when the initial frequency is small, *f*_0_ ≪ (*s*_*e*_*N*_*e*_ + *δ N*_*e*_*/*2)^−1^ (equation S37). In this limit, we see that the fixation probability can be approximately divided into two major regimes (Figure 7A; colored lines). For *small s*_*e*_, the fixation probability approaches a constant value. For *large s*_*e*_, decoupling noise becomes negligible compared to selection, and the fixation probability approaches the classical expression for fixation probability. The transition point between the two regimes is approximately,

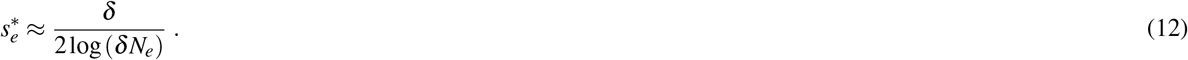

This transition point 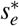 acts as a “decoupling-drift barrier,” below which selection cannot efficiently distinguish between genotypes with different fitness effects. This threshold can significantly exceed the classical “drift barrier” of 1*/N*_*e*_. In the strong decoupling/small initial frequency limit, we can express the fixation probability using an approximate piecewise function:

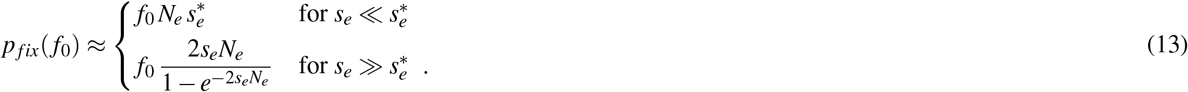

We compared our exact analytical expression for *p* _*fix*_ with simulations and the piecewise approximation (Figure 7A). The transition point 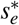 and piecewise approximation for *p* _*fix*_ show good agreement with both the exact expression and simulations. However, simulation results begin to diverge from the analytical expression at higher *δ*, as the first-order, small frequency deviation assumption in the Langevin equation becomes less accurate. Additionally, the *p* _*fix*_ approximation for 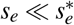 begins to deviate from simulations and analytics at higher *δ* due to the breakdown of the small *f*_0_ approximation.

Counterintuitively, we find that *p* _*fix*_(*f*_0_) *> f*_0_ for neutral or nearly-neutral mutants (i.e., mutants with *s* → 0). While one might expect *p* _*fix*_(*f*_0_) = *f*_0_ for neutral mutants due to symmetry, this is not the case when *δ* ≠ 0. Under these conditions, the mutant and wildtype are not exchangeable, as individuals from different genotypes covary differently. In fact, decoupling noise induces an effective frequency-dependent fitness effect (Figure S15; equations S12, S20), a phenomenon observed in similar models^21,48^. This effective frequency-dependent fitness effect is symmetric when *c*_1*A*_ = *c*_1*B*_. Consequently, a mutant experiencing decoupling noise but otherwise neutral with respect to the wildtype will have a disadvantage as it approaches fixation, but an effective advantage when rare–the crucial period for new mutants when they are most at risk for extinction. Thus, the increase in *p* _*fix*_ above *f*_0_ for neutral mutants directly results from this effective selective advantage for rare mutants.

### Site frequency spectrum

The site frequency spectrum (SFS) is a commonly used summary of the genetic diversity within a population. It describes the expected density of derived alleles at a given frequency; specifically *p*_*SFS*_(*f*)*d f* is the number of derived alleles in the frequency range [*f* − *d f /*2, *f* + *d f /*2]^60^. The population mutation rate is *θ*. Different dynamical processes can leave different characteristic signatures on the site frequency spectrum, so empirical site frequency spectra are often measured to infer aspects of the underlying evolutionary dynamics.

We calculate the SFS for alleles affected by decoupling noise by leveraging previously described approaches^61,62^. We find a general closed form solution for the SFS with constant selection and decoupling noise (SI section S3.2). We see that at low frequencies, the SFS both with and without decoupling noise decays as ∼ 1*/ f*, which is also the expectation for purely neutral alleles^38^ (Figure 7B). Simulations generally agree well with the analytics, but we see again that they start to deviate at higher *δ*.

A major feature of the SFS with decoupling noise lies in an uptick at high frequencies, which appears even in the absence of genuine (constant) selection (Figure 7B). Specifically, under strong decoupling noise *δ* ≫ 1*/N*_*e*_, and weak selection, we observe that the SFS approaches a constant when (1 − *f*) ≪ *δ N*_*e*_,

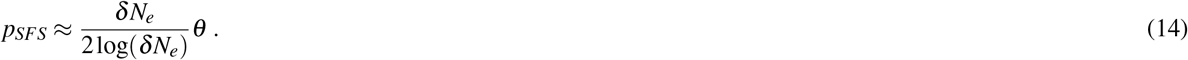

This is reminiscent of the classic SFS in the case of weak decoupling noise and strong selection, where the SFS also approaches a constant in the limit that (1 − *f*) ≪ (*N*_*e*_*s*_*e*_)^−1^,

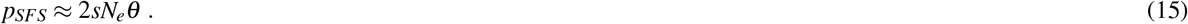

In the case of weak decoupling noise and strong selection, the SFS in the intermediate-high frequency regime (*N*_*e*_*s*_*e*_)^−1^ ≪ (1 − *f*) ≪ 1 scales like 1*/*(1 − *f*),

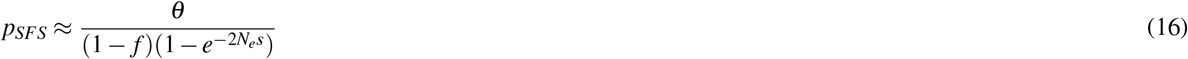

The SFS in the limit of strong decoupling noise and weak selection has a slightly different scaling behavior in the analogous regime, (*δ N*_*e*_)^−1^ ≪ (1 − *f*) ≪ 1, that includes a factor of log(1 − *f*)*/*(1 − *f*),

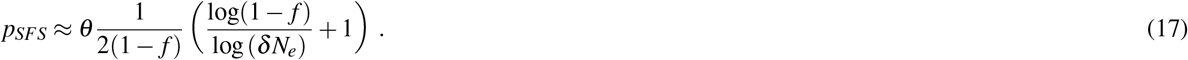

Decoupling noise can also substantially reduce diversity at intermediate frequencies. When *δ* ≪ 1*/N*_*e*_ and *N*_*e*_*s* ≫ 1, the minimum of the SFS is simply 4*θ*, at *f* = 0.5. In contrast, when *δ* ≫ 1*/N*_*e*_ the SFS at the same location is always lower,

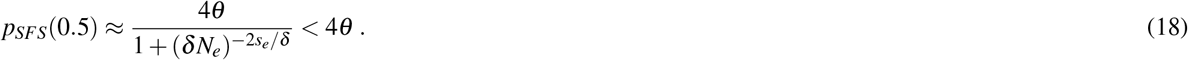

The value *p*_*SFS*_(0.5) ranges continuously from 4*θ* to 2*θ* as *δ* goes from zero to infinity. Overall, we showed how decoupling noise can decrease diversity at intermediate frequencies, while increasing diversity at high frequencies.

## Discussion

In this study, we show how an ecological mechanism can induce anomalous frequency fluctuations that are much larger than what can be attributed to classical genetic drift or measurement error. While classical genetic drift arises from independent offspring number fluctuations, our analysis suggests that these giant fluctuations arise when the offspring numbers of individuals are more correlated *within* genotypes compared to *between* genotypes. We constructed a simple and generic effective model of genotype frequency fluctuations, revealing that their magnitude is the sum of linearly-scaling classical genetic drift, and quadratically-scaling decoupling noise. By measuring the scaling relationships of the population fluctuations, we showed that the observed large fluctuations were indeed caused by decoupling noise.

Population abundance fluctuations are a well known, near-universal feature of populations across the tree of life, many of which follow Taylor’s power law^1–5^. Our effective model provides a general description of abundance fluctuations with few assumptions, valid for a range of underlying mechanistic processes. Fluctuating selection is one possible mechanism to explain such abundance fluctuations^21,48^, but importantly, other mechanisms such as chaotic dynamics^23,24^ and spatial effects^1,22^ can also cause such fluctuations. Importantly, our model predicts that any population composed of two or more genotypes that have sufficiently decorrelated offspring number fluctuations will experience decoupling noise. Many populations experience abundance fluctuations with a Taylor’s law exponent near two^19,20^, including unrelated experimental microbial populations^7^, and thus have correlated offspring number fluctuations. Decoupling frequency fluctuations are thus likely common, especially if offspring number correlations have a genetic basis and are mutable.

In our system, the correlated offspring numbers, and decoupling noise, appear to be caused by underlying chaotic dynamics. The chaotic dynamics produce a fluctuating selection-like effect, leading to mean-variance power law exponents of two^24^. Chaotic dynamics are known to be possible even in simple, single-genotype populations^25,63^. There are several plausible mechanisms that could generate chaotic population dynamics in our system, including overcompensation and nonlinear feedback loops between secreted metabolites and growth rates. The fluctuations appear even under the most tightly controlled environment we could create–among replicates from the same mother culture in the same shaking water bath. Thus, unlike fluctuating selection caused by a varying environment, we believe that this decoupling noise is inescapable, and an intrinsic aspect of the system. The underlying source of the decoupling noise in the unrelated barcoded *S. cerevisiae* populations that we reanalyzed^55,56,64^ remains unclear. As previously mentioned, a number of mechanisms are known to cause giant abundance fluctuations^2–5,21,23,24,48^. However, the recently reported extreme sensitivity of barcoded yeast populations to subtle variations in the environment^55^ is consistent with chaotic dynamics. Chaotic abundance dynamics have been suggested to be common among wild populations^33^, and have been demonstrated in a number of carefully controlled laboratory^26–29^ and field^30–32^ systems. Together, this opens the possibility that chaotic dynamics are also common in experimental microbial populations, and could play an important role in evolutionary dynamics by influencing genotype frequency dynamics across different systems.

The results of our study have implications for the inference of fitness effects in diverse biological populations. Among other possible mechanisms, subtle environmental fluctuations can be amplified by the chaotic dynamics, leading to significant extrinsic decoupling noise, which can cause batch correlations among replicates grown in the same conditions. These batch correlations may be mistaken for a genuine fitness effect, especially if the autocorrelation time of the environment is on a similar timescale as the evolution experiment. Thus, special care must be taken when designing evolution experiments to measure genotype fitness effects. For example, experimenters could perform biological replicates separately on different sets of days, or continuously measure environmental variables (e.g. temperature, humidity) over the time course to quantify the effect of environmental fluctuations. The accuracy of fitness inference procedures may be improved by explicitly modeling the effects of decoupling noise, along with classical genetic drift and measurement noise. Fitness inference methods that leverage temporal correlations between alleles^65^ must be handled with care, lest chaotic or spatial effects be confused with genuine (classical) selection.

Decoupling noise could cause a number of emergent effects on evolutionary dynamics. For example, we showed that the fixation probability of mutants can be drastically shifted by the presence of decoupling noise; thus, the fate of a mutant will not only depend on its fitness effect, but also on its decoupling parameters. The distribution of mutant fitness effects (DFE) is currently thought to largely shape the evolutionary dynamics of adapting populations^66,67^. But the DFE may (partially) lose its predictive power in the face of decoupling noise; we may need to consider a more general joint distribution, between fitness effects, drift effects, and decoupling parameters, especially if the joint distribution has non-trivial structure. For example, if the fitness effects of mutants do not correlate with their decoupling effects, or if decoupling effects are same between mutants, then which mutants become successful would be well predicted by considering the DFE alone. However, if both fitness effects and decoupling effects can vary between mutants, the group of mutants that reach high frequencies and eventually become fixed may differ from what would be anticipated based solely on the DFE. More broadly, the concept of a fitness landscape may need to be updated to a more general fitness-decoupling-drift landscape; evolutionary trajectories through such a landscape will likely differ compared to the case where the decoupling and drift effects are held constant. Significant effort has been devoted to measuring and characterizing the fitness effects of genotypes across systems, i.e. mean offspring numbers. But comparatively little effort has been placed in measuring the drift and decoupling effects of genotypes, i.e. offspring number variance and covariances, even though they can strongly shape evolutionary fates. We thus advocate for an effort to more routinely measure drift and decoupling effects alongside fitness effects.

In our theoretical analysis, we focused on the strong selection-weak mutation regime, where beneficial mutations rise and fix before another establishes. However, it is still unclear how decoupling noise would change the dynamics under other regimes, for example in the clonal interference/multiple mutations regime^68^. The evolutionary dynamics in the clonal interference regime may quickly become complicated, as one would have to account for the full offspring number covariance matrix between all genotypes present (SI sectionS2.3), rather than just a single decoupling parameter. Simplifying the covariance matrix by applying natural symmetries or random matrices may ease analysis, and could reveal complex behaviors. Additionally, broad-tail offspring number distributions can emerge out of a diverse array of growth processes^15,69,70^, but their effects on dynamics may change if such distributions are correlated across individuals. Both extensions are likely fruitful avenues for future work.

Overall, we presented experimental measurements for the scaling behavior of population fluctuations, and showed that we could explain them through a generic and extendable theoretical framework. We derived new theoretical results, showing how decoupling noise can impact evolutionary dynamics. We found that decoupling noise can arise quite generally, through a number of distinct mechanisms, so we believe that they may be common across diverse biological populations.

## Methods

### Growth conditions, media, and strains

All of the experiments presented here were performed in Davis Minimal Media (DM) base [5.36 g/L potassium phosphate (dibasic), 2g/L potassium phosphate (monobasic), 1g/L ammonium sulfate, 0.5g/L sodium citrate, 0.01% Magnesium sulfate, 0.0002% Thiamine HCl]. The media used in the LTEE and all experiments presented here is DM25, that is DM supplement with 25mg/L glucose. For coculture experiments, we first inoculated the desired strain into 1mL LB + 0.2% glucose + 20mM pyruvate. After overnight growth, we washed the culture 3 times in DM0 (DM without a carbon source added) by centrifuging it at 2500x*g* for 3 minutes, aspirating the supernatant, and resuspending in DM0. We transferred the washed culture 1:1000 into 1mL DM25 in a glass tube. Generally, we grew 1mL cultures in a glass 96 well plate (Thomas Scientific 6977B05). We then grew the culture for 24 hours at 37^°^C in a shaking incubator. The next day, we transferred the cultures 1:100 again into 1mL DM25. After another 24 hours of growth under the same conditions, we would mix selected cultures at desired frequencies, then transfer the mixture 1:100 to DM25. After another 24 hours of growth under the same conditions, we proceed with the experiment and start collecting measurements.

We used strains with fluorescent proteins inserted at the *attTn*7 locus, integrated via a miniTn7 transposon system, as previously reported^51^. The 6.5k *S* strain was tagged with eBFP2, the 6.5k *L* strain was tagged with sYFP2, REL606 was tagged with sYFP2, and the REL606 Δ*pykF* mutant^71^ was tagged with mScarlet-I.

### Flow cytometry

For all population measurements taken with flow cytometry, we used the ThermoFisher Attune Flow Cytometer (2017 model) at the UC Berkeley QB3 Cell and Tissue Analysis Facility (CTAF). For every measurement, we loaded the samples into a round bottom 96 well plate, for use with the autosampler. We set the flow cytometer to perform one washing and mixing cycle before each measurement, and ran 50 µL of bleach through the autosampler in between each measurement to ensure that there was no cross-contamination between wells. We used the “VL1” channel to detect eBFP2 fluorescence, which uses a 405nm laser and a 440/50nm bandpass emission filter. We used the “BL1” channel to detect sYFP2 fluorescence, which uses a 488nm laser and a 530/30nm bandpass emission filter. We used the “YL2” channel to detect mScarlet-I fluorescence, which uses a 561nm laser and a 620/15nm bandpass emission filter. We used a previously described and validated analysis framework^51^ to extract cell counts and strain frequencies from raw flow cytometry data. We used a previously described gating strategy, where we used threshold gates to separate clearly fluorescent particles from debris/noise (Figure S2 from Ascensao et al. (2024)^51^).

### Multi-day timecourses

We analyzed barcode sequencing data previously reported in Ascensao et al. (2023)^34^, focusing on experiment “Eco Eq 1”. We used the *N*_*e*_ estimate reported in Ascensao et al. (2023), which was obtained by decomposing within-ecotype neutral barcode frequency fluctuations into a component that accumulates over time (demographic fluctuations), and a component that is uncorrelated over time (measurement noise). We used previously reported data on colony forming units (CFUs) to estimate the bottleneck size, *N*_*b*_, by taking the average of barcoded (kanamycin resistant) CFUs over the timecourse. We obtained an estimate of the frequency variance between-ecotypes by fitting a linear model to the time course via ordinary least squares, and computing the variance around the line. We obtain quantitatively consistent results by leveraging the two biological replicates of the experiment, which were split at day 0, by calculating the variance between frequencies at day 1; this estimation method results in much wider confidence intervals, but a lower bound that is still several orders of magnitude larger than expected variance from bottlenecking and classical genetic drift. Similarly large between-ecotype fluctuations were found for all other coculture experiments presented in Ascensao et al. (2023).

We propagated cocultures of *S* and *L* (Figure1D-E), along with cocultures of REL606 and the REL606 Δ*pykF* mutant (FigureS2), where we started the cocultures as described above. We split the cocultures into eight replicates at day 0, where all replicates where grown in the same 37^°^C incubator, at the same time. We took flow cytometry measurements of the populations at the end of each 24 hour cycle. We computed a robust estimate of the variance (to decrease the influence of outliers) through the median absolute deviation, var *f* ≈ 2.1981 · (med_*i*_ |*f*_*i*_ − med_*i*_ *f*_*i*_ |)^2^. Confidence intervals were determined by standard bootstrapping. To compute the “genetic drift prediction” of how frequency variance should change over time, we used *N*_*e*_ = 10^5^, which is the approximate (conservative) bottleneck population size for both cocultures. We computed the approximate expected variance due to drift as var *f*_1_ = *f*_0_(1 − *f*_0_)*/N*_*e*_ for the first time point, and then var *f*_*t*+1_ = var *f*_*t*_ + ⟨ *f*_*t*_ ⟩ (1 − ⟨ *f*_*t*_ ⟩)*/N*_*e*_ for all subsequent time points.

To obtain estimates of the scaling behavior of variance quantities with respect to the frequency of the minor genotype, *S* (Figure2), we first set up *S*/*L* cocultures as described above, by mixing the cultures at different frequencies over about two orders of magnitude with *S* in the minority. We split each coculture into 16 replicate cultures, all grown under the same conditions. After one 24 hour growth cycle, we took three, independent flow cytometry measurements of each culture, which we treated as technical replicates for each culture. We utilized technical replicates for each culture primarily to decrease the effective amount of measurement noise for abundance–abundance measurements are noisier in our system than frequency measurements^51^. We took the average across technical replicates as the final frequency and abundance estimates for each biological replicate culture. We then computed variance across biological replicates with the standard estimator. Quantitatively and qualitatively similar results are obtained by using the robust estimator for the variance. We compared the variance to the mean frequency, instead of the frequency at the beginning of the cycle, because the within-cycle frequency dynamics show that the mean frequency is a better measure of the frequency right before variance starts to accumulate (Figure4A). We inferred the power-law exponents by performing ordinary least squares regression on the log-transformed variance/covariance quantity against the log-transformed mean *S* frequency; we determined confidence intervals via standard bootstrapping (Figure S3).

We measured the relationship between initial abundance and variance of abundance (Figure E8) by growing cultures as previously described, mixing *S* and *L* at around *f*_*S*_ ≈ 0.05 (the approximate equilibrium frequency) for the coculture condition, as well as continuing to propagate monocultures of *S* and *L*. We split the cultures into 16 replicate cultures per volume condition (using the same 1:100 daily dilution rate for all conditions). We used 0.1, 1, and 10mL culture volumes, where used the glass 96 well plates for the first two conditions, and 50mL glass erlenmeyer flasks for the 10mL condition. After the 24 hour growth cycle, we plated each replicate culture on LB plates, at a 10^−5^mL^−1^ dilution rate. We additionally took flow cytometry measurements for each replicate in the coculture condition, to more accurately measure genotype frequency. Confidence intervals were computed by standard bootstrapping. There are no statistically significant differences in variance scaling when comparing the monoculture and coculture conditions, or between *S* amd *L*.

### Within-cycle timecourses

After the initial growth cycles of fluorescently tagged *S* and *L* as previously described, we mixed the strains together such that the relative frequency of *S* was around 6%. We grew the coculture for one more cycle in DM25, then took a flow cytometry measurement at the end of the 24 hour cycle, which we took as time 0. We then immediately inoculated new replicate cultures from the overnight mother culture by diluting the culture 1:100 into DM25 (e.g. 300 µL of culture + 30mL of DM25), vortexing the mixture well, and then splitting the resulting mixture into 1mL cultures in individual wells of a glass 96 well plate. We used 5 biological replicates for the 8 hour time-course and 23 replicates for the 24 hour time-course. We secured the 96 well plate in a 37^°^C water bath, shaking at 180rpm. The wells of the glass 96 well plate are separated such that water can pass in between the wells. We briefly removed the plate at designated time intervals (about every 30 minutes and 2 hours for the 8 and 24 hour time course, respectively) to subsample approximately 60 µL of culture for flow cytometry measurements. Subsamples were discarded after measurement. Exact times of plate sampling were documented. We subsampled the cultures from the 24 hour time course in two batches (one of 11, one of 12), where we subsampled the second batch immediately after the flow cytometry measurement of the first batch was finished. We utilized the batch structure to minimize the amount of time the samples have to wait outside of the water bath to be measured by the flow cytometer.

Following data processing, we computed the variance between biological replicates at the same time points. For the 24 hour time-course, we first computed the variance among all samples in the same batch for each time point, then averaged the variance between the two batches at the same time point, because the batches were taken at slightly different actual times. The confidence intervals for variance measurements were computed with standard asymptotic formulas. We fit various curves (Figure S7A) to the variance trajectory after 7 hours by first log-transforming the variances (as an approximate variance-stabilizing transform), and then performing least squares regression. Denoting *v*_*i*_ as the variance and *t*_*i*_ as the time, we fit a simple exponential curve, log *v*_*i*_ = log *a* + *bt*_*i*_ + *ε*_*i*_, a linear curve, log *v*_*i*_ = log(*a* + *bt*_*i*_) + *ε*_*i*_, a generalized power-law curve, 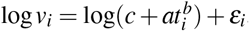, and a quadratic curve, log 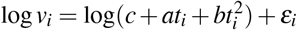 Confidence intervals for the fits were obtained by resampling trajectories of biological replicates with replacement (standard bootstrapping), computing the variance between replicates in the same manner as previously described, and performing the appropriate regression again.

We evaluated the fits of all regressions by computing their Akaike Information Criterion (AIC),

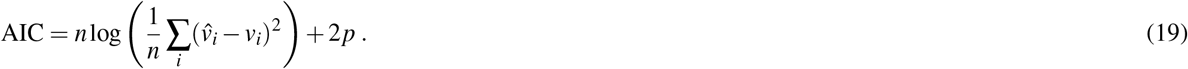

Where 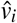 is the variance predicted from the regression model, *n* is the number of time points, and *p* is the number of parameters fit in the model. We observed that the AIC of the exponential fit was lower than all others (Figure S7B). Thus, we sought to determine if this difference was significant. To this end, we computed paired one-sided p-values similarly to how we computed confidence intervals, i.e. with standard bootstapping, re-performing the regression, and calculating the AIC.

We calculated Lyapunov exponents (*λ*) from the frequency trajectories after seven hours in two ways. First, we implemented and appropriately modified the method proposed by Rosenstein et al. (1993)^53^, based on nearest-neighbor distance (NND) trajectories. The method relies on (1) appropriately embedding the data, (2) identifying the nearest neighbor *j* at the initial timepoint of each trajectory *j* (calculated using euclidean distance), (3) computing the euclidean distance between initially nearest neighbors over time, *d*_*i, j*_(*t*), and (4) fitting an exponential to all computed pairwise distances via least squares regression,

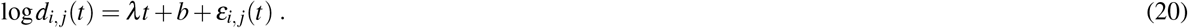

To embed the data, we first needed to choose the appropriate hyperparameters for the embeddding dimension and lag time, which we did via shuffle splitting cross-validation. Specifically, for each embedding, we shuffled the data with 1000 replicates, then used 30 of the distance points to fit the model shown in equation 20 and obtain and estimate for *λ* and *b*. Using the fitted parameters, we computed the mean squared error (MSE) for all remaining out-of-sample points, and averaged the MSE across all shuffled replicates. The one-dimensional case had the lowest out-of-sample MSE, so we proceeded with the one-dimensional Lyapunov exponent estimate (Figure S8B). The Lyapunov exponents obtained for the other hyperparameter sets were very similar to the one-dimensional case (Figure S8A), showcasing the robustness of the method.

Recent work has suggested the use of Lyapunov exponent inference methods based on dynamics reconstruction and jacobian estimation^33^. However, our timecourse is not long enough to accurately reconstruct dynamics in the necessary manner, and would not allow for effective jacobian estimation. Additionally, measurement noise is small in our experimental set-up, and we can leverage the numerous biological replicates starting from (nearly) the same initial conditions to directly look at divergence of trajectories. Thus, we do not believe the use of a jacobian-based inference method is necessary or appropriate for this experiment.

The exponentially increasing variance between biological replicates can also be used to extract a Lyapunov exponent (SI section S4.1). Specifically, var *f* ∝ *e*^2*λt*^, so we took the Lyapunov exponent as half of the fitted exponent. This is valid if the system is well-described as one-dimensional (although this assumption could be relaxed in principle); our previous hyperparameter analysis revealed that this was indeed the case. In both methods, we obtained confidence intervals by standard bootstrapping.

### Extrinsic fluctuations and splitting cultures

We started a coculture of *S* and *L* with the same protocol as previously described. At day 0, we split the culture into 4 replicate cultures by diluting the culture 1:100 into DM25, vortexing the mixture well, and then splitting the resulting mixture into 1mL cultures in individual wells of a glass 96 well plate. We split replicate cultures at days 3 and 7, via the same procedure, into 3 and 2 new subreplicates for each culture respectively. We took flow cytometry measurements at the end of each growth cycle for twelve days.

We sought to model the effects of measurement noise, frequency-dependent fitness effects, intrinsic and extrinsic decoupling noise, and memory-like effects from sharing mother cultures (from splitting cultures). We used a Bayesian hierarchical modeling approach to model the data, as it allowed us to flexibly set up the model, and obtain full posterior estimates. We focused on modeling the logit frequency displacements for each culture, because a logit transform serves as a variance-stabilizing transform for decoupling noise and fitness effects. For example, in the simplest case, considering just fitness effects and intrinsic decoupling noise, the distribution of frequencies at time *t* is,

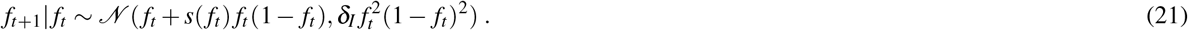

After a logit transformation, the frequency displacement is,

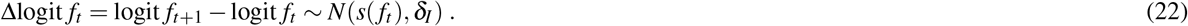

We model the effect of environmental fluctuations by considering an environmental effect that is drawn from a centered normal distribution at each time point,

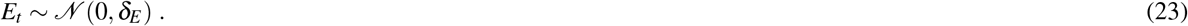

Similarly, we model the effect of shared mothers as another centered normal distribution,

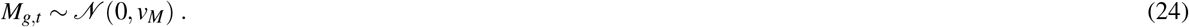

The *g* subscript indexes a group that all arise the same mother culture at time *t*, such that all daughter cultures will share the same *M*_*g,t*_. We focus on modeling the effect of sharing a mother culture immediately after splitting cultures on days 3 and 7; for all other days, we set *M*_*g,t*_ = 0. Both the effect of shared mothers and environmental fluctuations feed into the mean of the frequency displacement for each timepoint. We model the frequency dependent fitness as a linear function, *α* + *β f*_*t,g,i*_, which appears to capture this dependence well in the frequency regime our data lies in (Figure S10). The final hierarchical model that we fit to the data reads as,

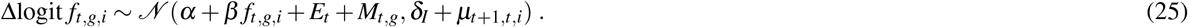

We account for measurement noise by specifying *µ*_*t*+1,*t,i*_; we previously found that frequency measurements have errors well approximated by a binomial distribution in our flow cytometry protocol and set-up^51^. For the logit-transformed frequency displacements, we use a first order approximation for the binomial variance such that,

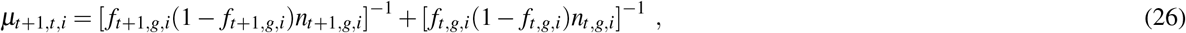

where *n*_*t,g,i*_ is the total number of cells detected in the flow cytometer.

We fit the model to all of the data (frequency displacements), jointly inferring *α, β, δ*_*I*_, *E*_*t*_, *M*_*t,g*_, *δ*_*E*_, and *v*_*M*_. We use the Hamiltonian Monte Carlo (HMC) algorithm implemented in STAN^72^ to jointly estimate the posterior of each parameter. We used a non-centered parameterization of the model to improve convergence of the hierarchical model. For the variance parameters, i.e. *δ*_*E*_, *δ*_*I*_, and *v*_*M*_, we use the Jeffrey’s prior for variance, *p*(*x*) ∝ *x*^−2^. To account for the non-centered parameterization and *Mt,g*, we use a standard normal prior, *x ∼ N*(0, 1), then multiplied both by the standard deviation parameters, 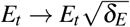 and 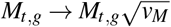 For all remaining parameters, we used a uniform prior.

### Reanalysis of barcoded yeast data

We reanalyzed the barcoded yeast strain time-courses from Venkataram et al. (2016)^56^. We excluded batch 2 from our analyses because it only had two biological replicates, while batches 1, 3, and 4 all had three biological replicates each. The serial dilution evolution experiments in Venkataram et al. were conducted such that there was one culture at time point 1, which was sampled for barcode sequencing, which was then split into three (or two) biological replicates, which were all sampled for all subsequent time points. We first computed the mean and variance of each barcode frequency across the three biological replicates at time point 2 (after one growth cycle apart). We only pooled mean-variance data points from batches 1 and 3 in Figure 6A because batch 4 behaved slightly differently than the other two, in that it had stronger decoupling noise (Figure 6B,D). However, the data from batch 4 still has the same asymptotic scaling behaviors (Figure S11). After plotting the relationship between the mean and variance of barcode frequencies, we computed a moving average of the relationship. We computed the moving average of var(*f*) as a function of ⟨ *f* ⟩with a multiplicative window (i.e. a constant sized window on log ⟨ *f* ⟩). We went along the x-axis, and computed the average of var(*f*) in between [⟨ *f* ⟩ · (1 ™ *w*), ⟨*f* ⟩ · (1 + *w*)], where we set *w* = 0.3. Changing the smoothing parameter *w* does not significantly alter the results. We computed confidence intervals by standard bootstrapping. We estimated power law exponents by performing ordinary least squares regression on the log-transformed variance against the log-transformed mean. For the low-frequency power-law fit, we used all points below *f <* 5 10^−5^. For the high-frequency power-law fit, we used all points above⟨ *f* ⟩ *>* 3 10^−4^. We chose those frequency ranges because they represented the approximate maximum ranges where the apparent power-law slope does not deviate significantly from the “asymptotic” power-law slope near the boundaries of the frequency ranges. We computed standard errors on the power-law exponents by standard bootstrapping.

We then sought to estimate the mean squared displacement (MSD) of the log-transformed frequencies, at high frequencies such that decoupling noise is dominant (Figure 6B). The log transformation acts a variance-stabilizing transform; we can use this approximation (instead of a logit transform) as all barcodes here are at low frequency *f* ≪1. We expect that the MSD should linearly increase over time increments, with a slope determined by *δ*,

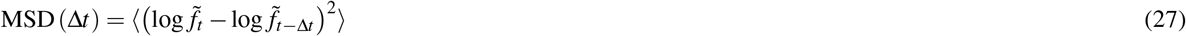

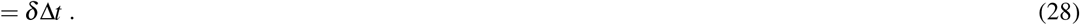

We estimated the MSD for all pairs of time points that were either 1 or 2 time increments (Δ*t*) apart. We filtered for high frequency barcodes at a mean frequency range from 5 · 10^−4^ to 5 · 10^−3^. We took 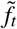 as centered barcode frequencies, where we subtracted the average barcode frequency over all replicates in a batch, 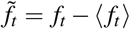. This is equivalent to subtracting the mean “fitness effect” over a pair of time points. We need a way to separate measurement error (uncorrelated in time) from biological decoupling noise (accumulates over time). We leveraged a previously utilized^34^ method to pull apart the two sources of noise. Briefly, in the presence of measurement noise, variance between two time points will be the sum of the decoupling noise and measurement noise (ζ_*t*_) for the two time points,

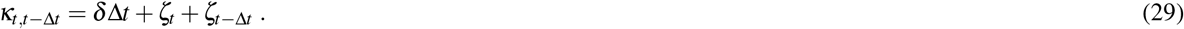

For every barcode, we compute the log difference at considered time point pairs, 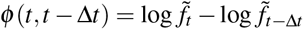. We then obtain a robust estimate of *κ* through the median absolute deviation, over all barcodes,

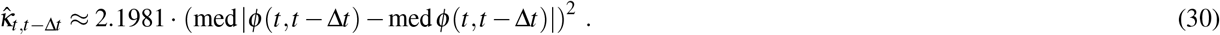

We obtain estimates for the standard error, std 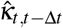, through standard bootstrapping. With the relationship between *κ*_*t,t*−Δ*t*_ and the noise parameters (equation 29), we can estimate the noise parameters given all of our measured 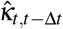. We do this by numerically minimizing the weighted squared difference, 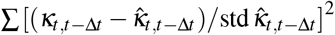. We obtain our final estimate of the MSD by subtracting the measurement noise parameters from each 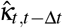, and averaging over all values with the same time increment Δ*t*,

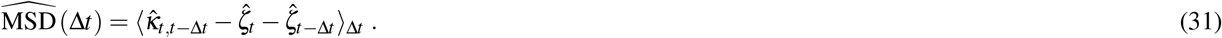

We obtained confidence intervals for the MSD via standard bootstrapping. We are uncertain about the scaling behavior of the measurement noise parameters, relative to the mean frequency. We treat the ζ_*t*_ parameters as constant, which should be a valid assumption if the measurement noise does not significantly change over the frequency range that we used. Using an even smaller frequency range does not seem to change our results significantly–we obtain similar results with a thinner mean frequency range of 7 · 10^−4^ to 2 · 10^−3^, albeit with higher error (as expected) (Figure S12).

To model the contributions of intrinsic/extrinsic decoupling noise to the frequency trajectories of high-frequency barcodes, we turned to a similar set-up as previously considered (equation 25). We change the model to reflect the data structure,

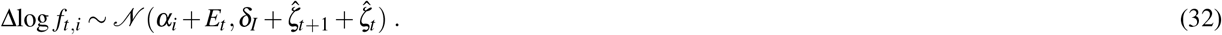

We eliminated the “sharing mothers” parameters, and we included a barcoded-dependent mean fitness effect *α*_*i*_, which we fit for each barcode. We fit the model for each batch in the same manner as previously described.

### Evolutionary dynamics simulations

We simulated the evolutionary dynamics of a two-genotype system to compare with our theoretical results on the evolutionary implications of decoupling noise, as shown in Figure 7. We directly simulated the number dynamics of each genotype, *N*_*i*_, with the Langevin equation (following the approach of Melbinger and Vergassola (2015)^21^),

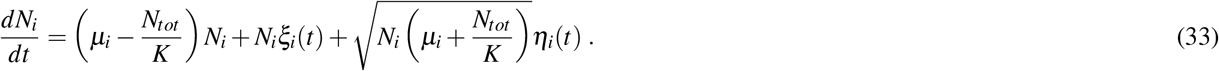

Here, *N*_*tot*_ = ∑_*i*_ *N*_*i*_, *η*_*i*_(*t*) is standard Gaussian white noise, and *ξ*_*i*_(*t*) is correlated Gaussian white noise, such that,

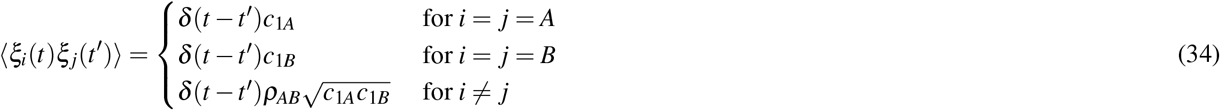

The frequency dynamics, *f* = *N*_*A*_*/*(*N*_*A*_ + *N*_*B*_), have previously been approximately derived from the abundance dynamics^21^, using a first order approximation. We see that those frequency dynamics are equivalent to the dynamics we derive (equation S20) when the classical genetic drift parameters are the same between genotypes, *κ*_*A*_ = *κ*_*B*_. The fitness effect becomes *s* = *µ*_1_ ™ *µ*_2_, and the effective population size becomes *N*_*e*_ = *K/*2. We directly simulated the abundance dynamics in equation 33 by discretizing time,

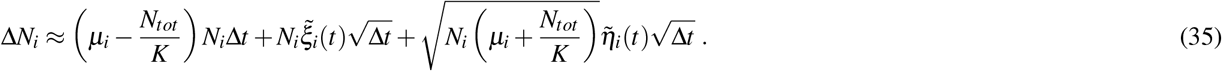

Now 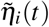 is a standard Gaussian random variable, and 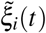 is a Gaussian random variable with mean of 0 and a covariance matrix given by equation 34. We used initial conditions of *N*_*A*_ = 10^−3^*K* and *N*_*B*_ = *K*. In our simulations, we consistently used *K/*2 = 10^3^, *µ*_*A*_ = 1, *ρ*_*AB*_ = 0.5, and *c*_1_ = *c*_1*A*_ = *c*_1*B*_. We define the correlation between A and B as 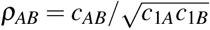. We set Δ*t* = 0.1, which seemed to sufficiently mitigate time-discretization error. We varied *c*_1_ and *µ*_*B*_ = 1 − *s* in our simulations.

For the simulations to estimate fixation probability, we ran 10^5^ independent simulations of the dynamics for each parameter set, where we ran the simulations until one of the genotypes fixed. We considered the major genotype as fixed when the minor genotype dropped below 0.1 in the population. We computed the fixation probability as simply the proportion of simulations where genotype *A* fixed in the population. Similarly for the simulations to estimate the site frequency spectra, we ran 2 10^4^ independent simulations of the dynamics for each parameter set. We recorded the genotype frequency at each timestep until one of the genotypes fixed, and the appropriately normalized the data to obtain the expected density of *A* at a given frequency bin.

## Code, data, and strain availability

All code and data presented in this manuscript are available athttps://github.com/joaoascensao/giantpopflucts. All strains presented in this paper are available upon request.

## Acknowledgements

We thank Adam Arkin, Kira Buttrey, Benjamin Good, Jonas Denk, Kelly Wetmore, QinQin Yu, Jimmie Ye, Dipti Nayak, and all members of the Hallatschek lab (past and present) for helpful comments and advice on the project. We thank Richard Lenski for sending us the LTEE-derived strains and populations, along with experimental advice and feedback. We thank Tim Cooper for sending us the REL606 Δ*pykF* mutant. Research reported in this publication was supported by a National Science Foundation CAREER Award (1555330). This work was supported by the National Institute of General Medical Sciences of the NIH under award R01GM115851 and by a Humboldt Professorship of the Alexander von Humboldt Foundation. JAA acknowledges support from an NSF graduate research fellowship, a Berkeley fellowship (from UC Berkeley), and Lloyd and Brodie scholarships (from UC Berkeley Dept of Bioengineering). We thank Mary West of the Cell and Tissue Analysis Facility (CTAF) at UC Berkeley. This work was performed in part in the QB3 CTAF, that provided the ThermoFisher Attune Flow Cytometer (2017 model).

---

**Extended Data Figure 8.**
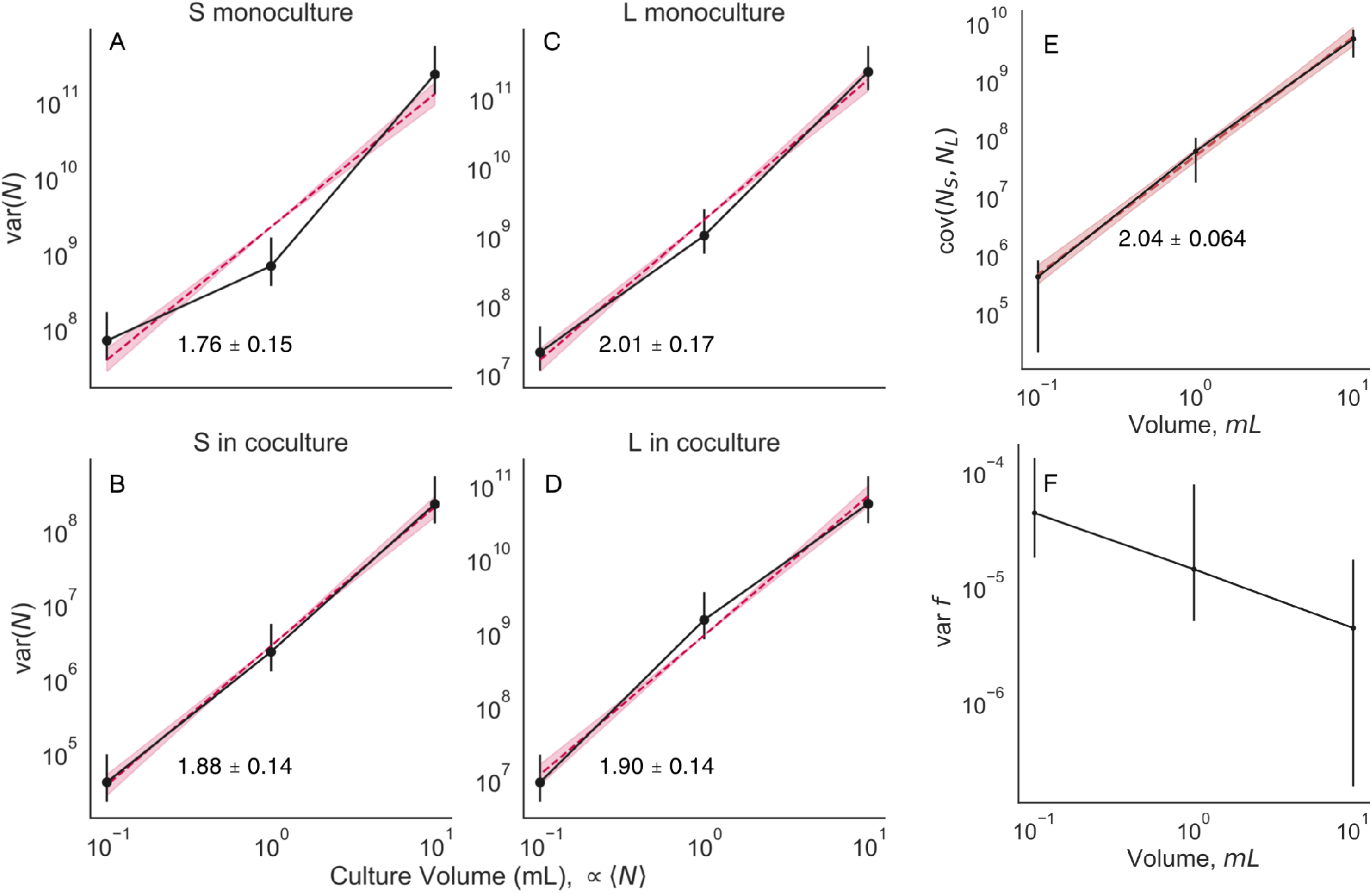
Relationship between abundance mean and variance (related to Figure 2). We cultured monocultures of *S* and *L*, along with a coculture of *S*/*L* at three different culture sizes (although with the same media, and at the same daily dilution rate). We split the cultures in each condition into 16 biological replicates, then measured abundance and coculture genotype frequency after one growth cycle. (**A-D**) The abundance variance scales with the mean like a power law, with an exponent of approximately 2. The exponent is not significantly different between genotypes, or between the monoculture/coculture condition. (**E**) The covariance between *S* and *L* abundance also scales like a power law with an exponent of two. (**F**) The frequency variance appears to decrease as a function of culture volume. This may represent evidence that the culture environments differ enough to cause changes in fluctuation strength. All error bars represent 95% CIs.

**Extended Data Figure 9.**
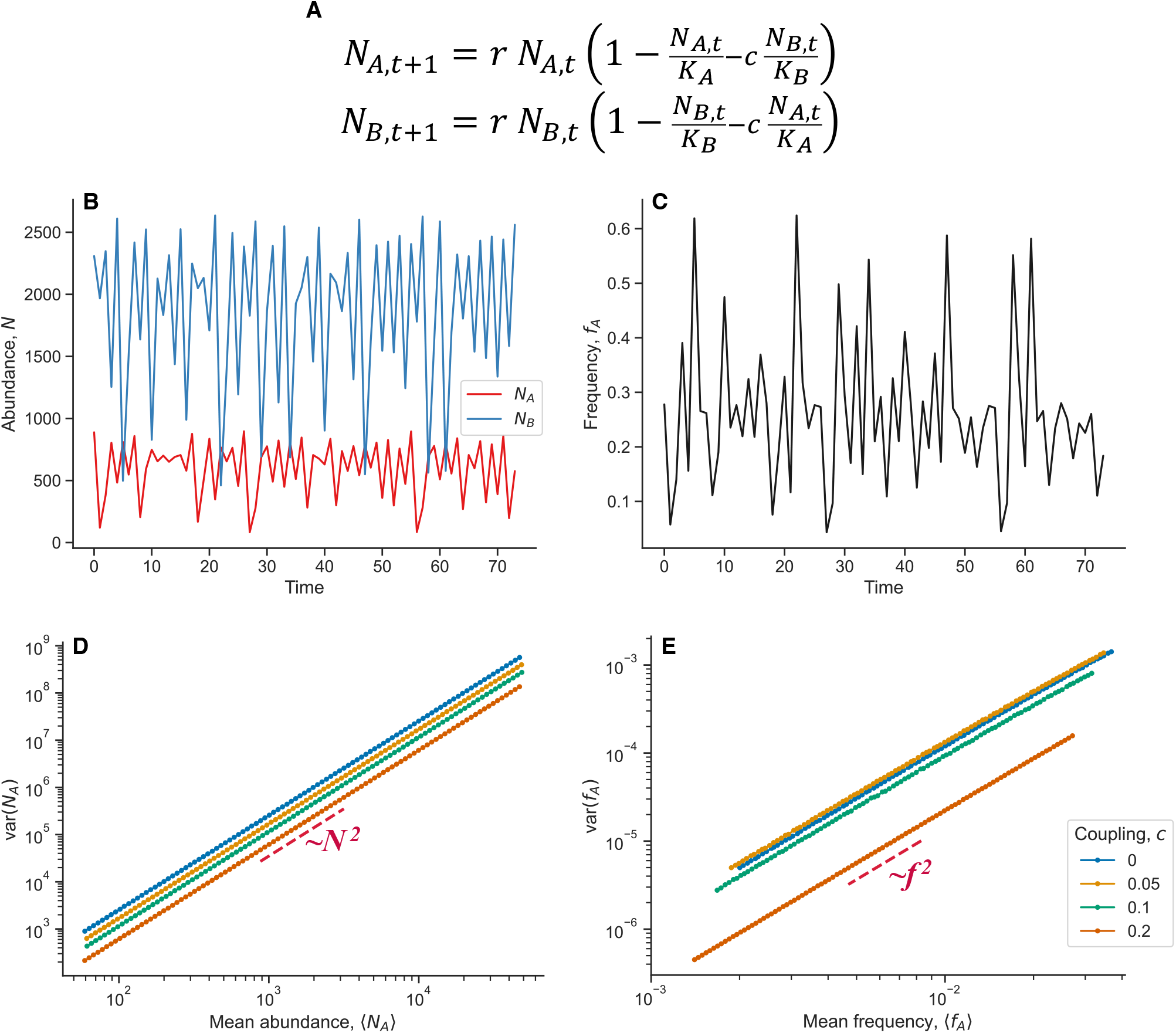
Simulations of chaotic dynamics in a simple two-genotype system (related to Figure 4). (**A**) We model the dynamics of two genotypes, *A* and *B*, as coupled logistic maps (in discrete time). The parameters *K*_*A*_ and *K*_*B*_ are the carrying capacities of strains *A* and *B*, respectively. When the coupling parameter, *c*, goes to zero, we recover standard, independent logistic maps. Here, we consistently use *r* = 3.9, which puts the populations in the chaotic regime. (**B-C**) Examples of the abundance and genotype frequency dynamics. Here, we use *c* = 0.1. Even though the abundance dynamics are mildly coupled to each other, we still see fluctuations creeping into the genotype frequency dynamics. (**D-E**) Mean-variance scaling behaviors of the population abundance and genotype frequency. In both cases, we see power-law scaling with exponents of two, indicating the presence of effective offspring number correlations and decoupling noise. We varied the carrying capacities over several orders of magnitude to change the abundance and frequency. We computed the mean and variance of trajectories ran for 10^5^ iterations, and discarded the first 10^4^ iterations (to control for transient behaviors). We see that the degree of coupling does not affect the power-law exponent, but can change the intercept (as expected).

**Extended Data Figure 10.**
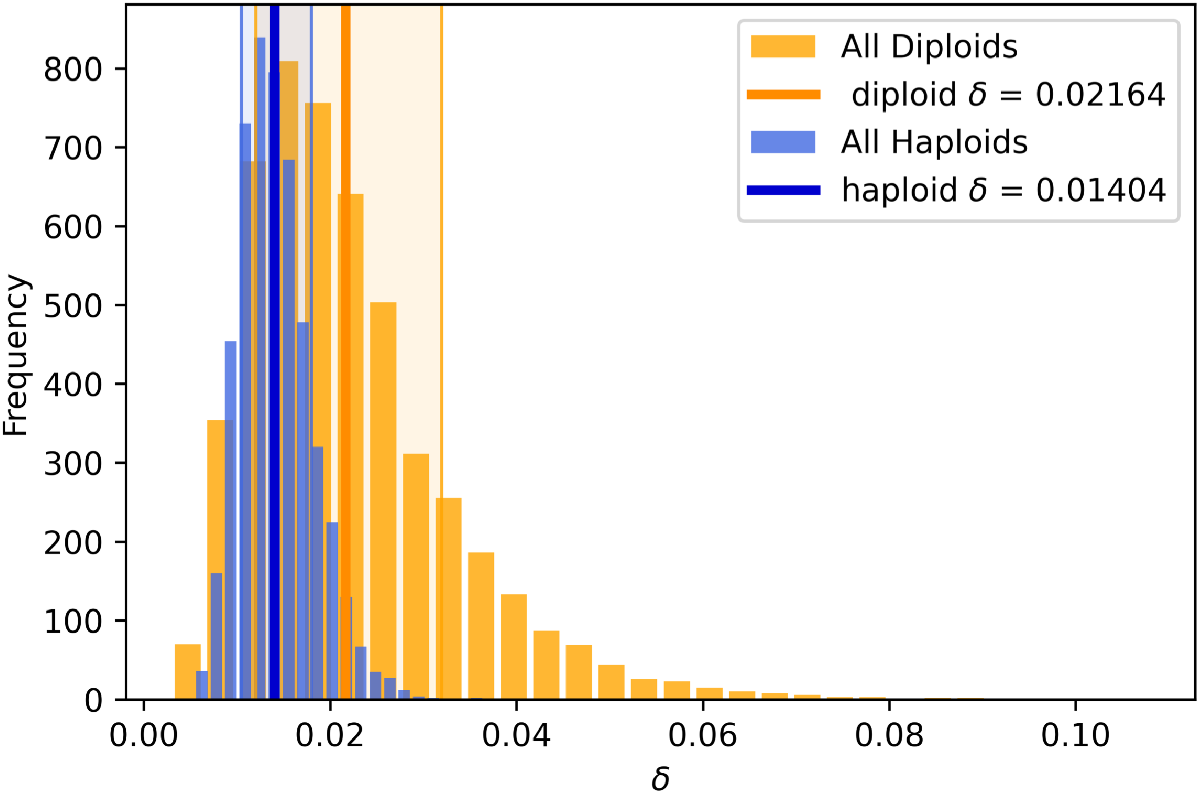
Estimate of sampling distribution of *δ*_*I*_ in Venkataram et al.^56^ data (related to Figure 6). We used standard bootstrapping to separately estimate *δ*_*I*_ for adaptive haploids and diploids. We find that there are not significant differences in the decoupling parameter between the two classes of beneficial genotypes. The solid lines represent means of the sampling distribution; shaded regions represent 68% confidence intervals.

**Extended Data Figure 11.**
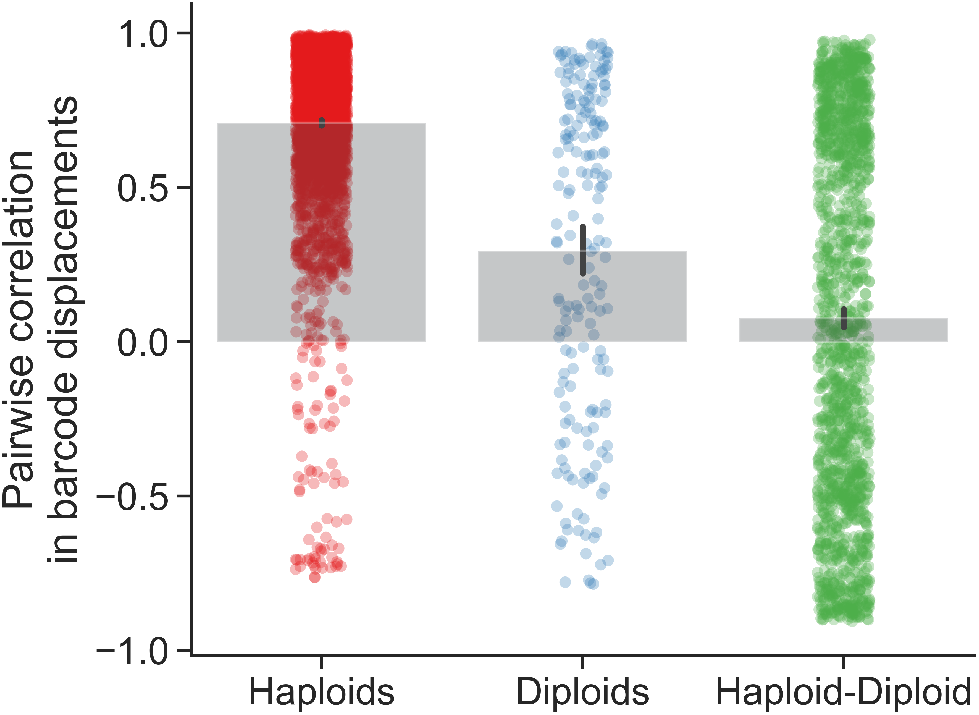
Correlated fluctuations between barcoded clones, from Venkataram et al.^56^ data (related to Figure 6). We computed the pairwise correlation in log-displacement between every pair of high-frequency clones over every time point, replicate, and batch, i.e. corr(Δlog *f*_*i,t*_, Δlog *f* _*j,t*_). Points represent correlation coefficients for every pair of clones, and bars represent the average. Error bars represent 95% CIs. We see that, on average, pairs of haploid clones have highly correlated displacements, followed by pairs of diploid clones, and then pairs consisting of one haploid and one diploid clone.

## Supplementary Information

## S1 Table of strains

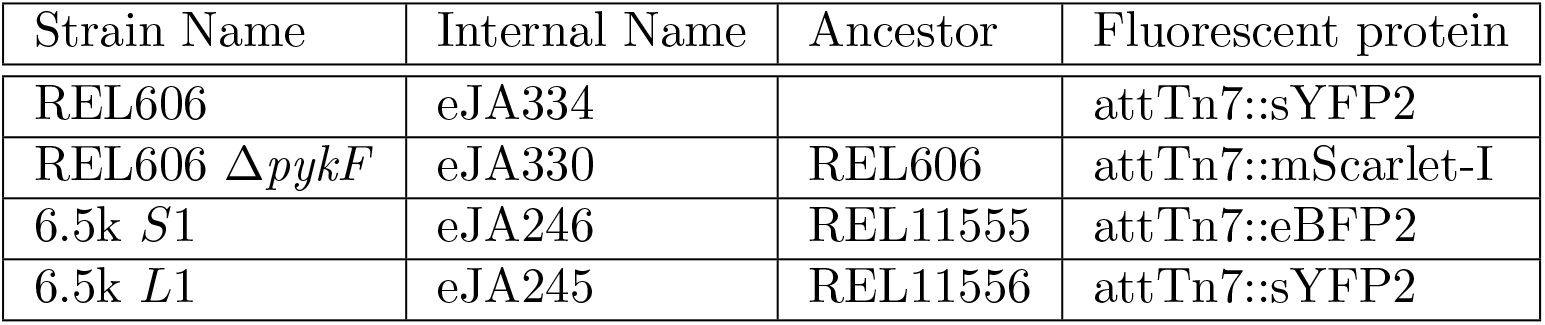

| Strain Name | Internal Name | Ancestor | Fluorescent protein |
| --- | --- | --- | --- |
| REL606 | eJA334 |  | attTn7::sYFP2 |
| REL606 $\Delta pykF$ | eJA330 | REL606 | attTn7::mScarlet-I |
| 6.5k *S1* | eJA246 | REL11555 | attTn7::eBFP2 |
| 6.5k *L1* | eJA245 | REL11556 | attTn7::sYFP2 |

## S2 Effective model of population fluctuations

### S2.1 Two-genotype case

To model the process of population fluctuations, we first consider a simple population consisting of one genotype, where all individuals are identical. The change in population size of a genotype *µ* from size *N*_*µ*_ in the current time point to size 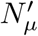 in the next time point can be decomposed as

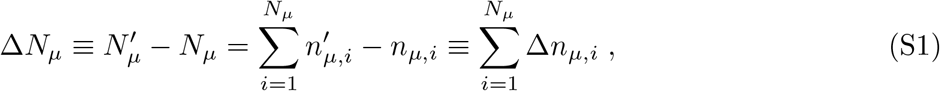

where Δ*n*_*µ,i*_ ∈ [− 1, 0, 1, …], *i* ∈ [1, *N*_*µ*_] is −1 plus the offspring number of the *i*^*th*^ cell of the *N*_*µ*_ cells of the *µ* genotype that exist at the current time step. We know that if all cells behave the same way, we have to demand

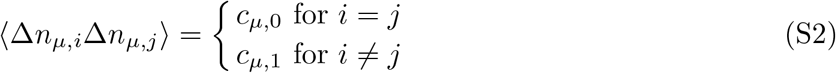

Here, the brackets ⟨·⟩ represent the mean of a random variable. This is the only mathematical form of the covariance parameters that won’t change if we relabel the cells–a symmetry argument that can easily be extended to multiple genotypes. This assumes that the offspring number distributions have finite covariances, i.e. are not heavy-tailed. Note that *c*_1_ *< c*_0_ follows from 0 *<* ⟨(Δ*n*_*i*_ − Δ*n*_*j*_)^2^⟩ = 2*c*_0_ − 2*c*_1_.

We can express the variance in total population size in terms of *c*_0_ and *c*_1_,

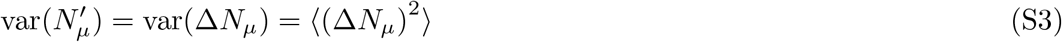

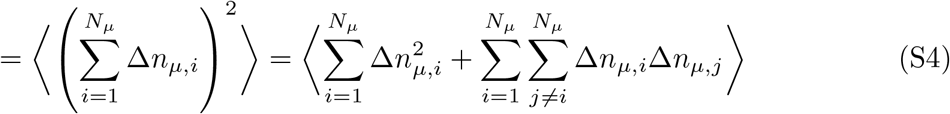

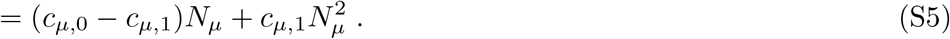

The form of this variance scaling has been previously noted [1, 2]. Power law mean-variance scaling of population abundance has been widely observed in ecology, where it is known as Taylor’s power law [3, 4, 5, 6, 7]. For large enough values of *N*_*µ*_, the distribution of 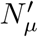 will converge in distribution to a gaussian random variable, because of the central limit theorem. So the fluctuations of 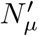 will solely depend on the variance, var 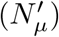.

We can now extend the analysis to a population consisting of two genotypes labeled *A* and *B*, possibly with non-identical properties (Figure 2A). We can generalize the below analysis to multiple genotypes (SI section S2.3). Extending the symmetry argument from above, we now require five covariance parameters to fully describe the population,

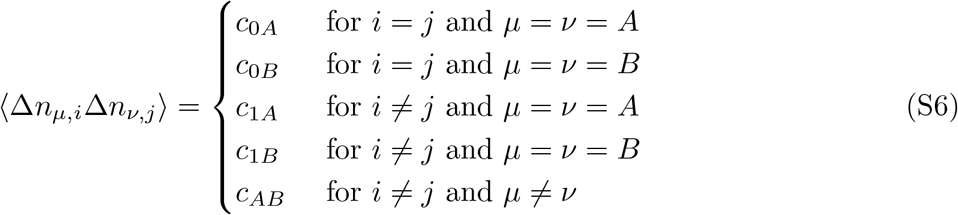

The expression for the variance of the total population size for each genotype is the same as in equation S5. The covariance of the total population sizes between genotypes is,

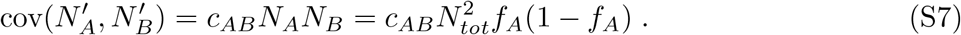

Where *N*_*tot*_ = *N*_*A*_ + *N*_*B*_ is the total population size, and *f*_*A*_ = *N*_*A*_*/*(*N*_*A*_ + *N*_*B*_) is the frequency of genotype *A* in the population. The frequency of a genotype will also change from one time point to the next, induced by the fluctuations of individuals. The expected frequency variance at the next time point can be written as,

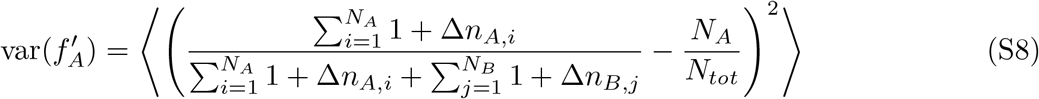

If we assume that the frequency deviations are small, we can expand in the deviations to obtain,

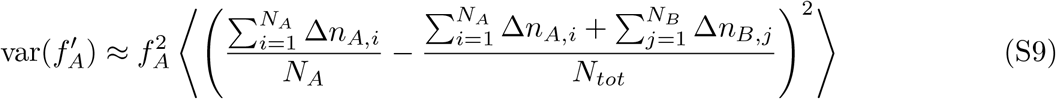

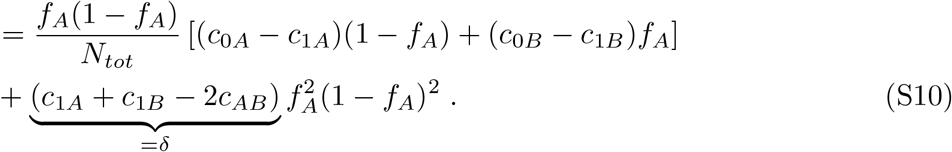

We can approximate the expected frequency by expanding to second order,

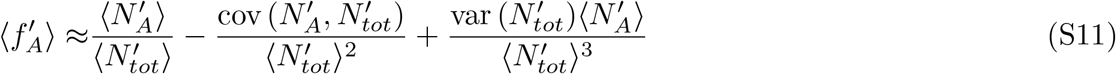

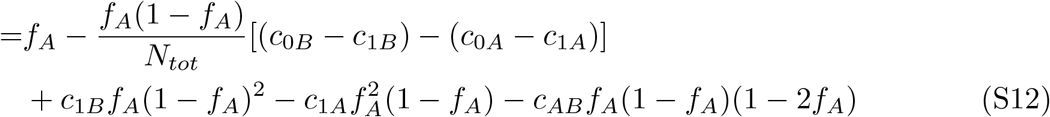

### S2.2 Fluctuating environments

We can incorporate the effects of a fluctuating environment into our model, decomposing offspring number fluctuations into intrinsic and extrinsic fluctuations. We consider a fluctuating environment that causes genotype fitness to fluctuate with gaussian noise, *w*(*E*) ∼ 𝒩(*s*_0_, *v*_*E*_). Conditioned on the environmental state, the covariance matrix ⟨Δ*n*_*µ,i*_Δ*n*_*ν,j*_ |*E*⟩ corresponds to equation S6, representing the intrinsic fluctuations of the system. And so the total abundance variance conditioned on the environmental state, var 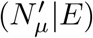, also corresponds to equation S5.

To consider the combined effect of the intrinsic and extrinsic fluctuations on the total population abundance, we must marginalize out the effect of environmental fluctuations,

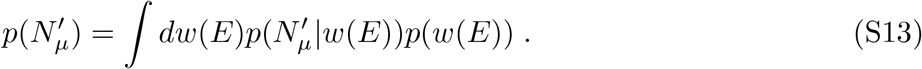

The conditional abundance density will be gaussian by the central limit theorem,

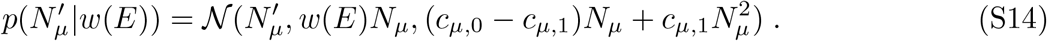

Here, 𝒩 (*x, µ, v*) is a standard gaussian density. We use a standard identity for the product of gaussian densities,

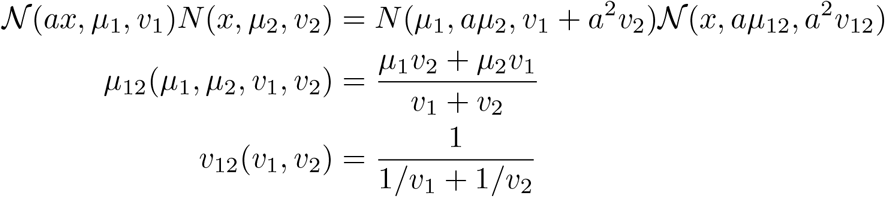

By rearranging, applying the above identity, and integrating equation S13, we obtain a final marginal density of

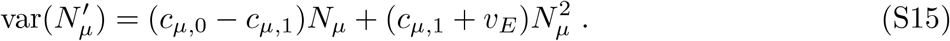

Thus, the total strength of correlated abundance fluctuations is simply the sum of the strength of fitness fluctuations (extrinsic source) and the strength of the intrinsic fluctuations. Applying the same logic, it immediately follows that the decoupling parameter *δ* can also be decomposed into intrinsic and extrinsic components, *δ* → *δ*_*I*_ + *δ*_*E*_.

### S2.3 Generalization to *m* genotypes

When there are *m* distinct genotypes/species present in a population, possibly with differing offspring number covariances, what is the expected frequency variance?

We first consider the possible covariance parameters between the change in offspring number for a single individual across a single generation, Δ*n*_*µ,i*_, where *µ* labels the genotype and *i* labels the individual. Again, Δ*n*_*µ,i*_ ∈ [−1, 0, 1, …], *i* ∈ [1, *N*_*µ*_] is −1 plus the offspring number of the *i*^*th*^ cell of the *N*_*µ*_ cells of the *µ* genotype that exist at the current time step. For *m* genotypes, we require 2*m* + *m*(*m* − 1)*/*2 covariance parameters:

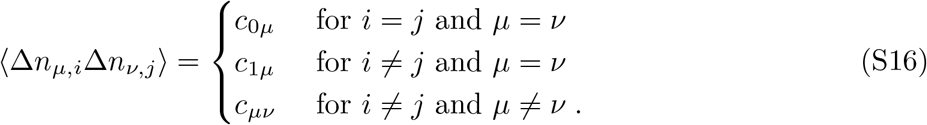

We define the initial frequency of genotype A (arbitrarily labeled) as,

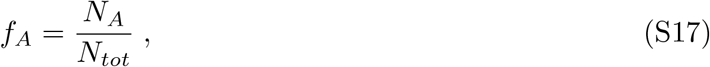

where 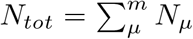. The frequency after one generation is similarly defined, and labeled as 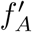. If we assume that frequency deviations are small, we can expand in the deviations to obtain,

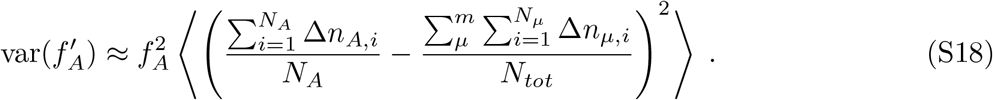

Following some algebra, and using equation S16, we arrive at the following expression,

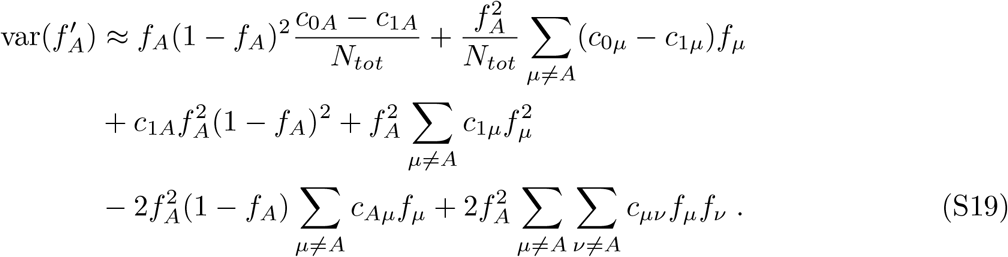

We can see that the generalized equation has a similar form to the two-species case, where the first two terms arise from independent offspring number fluctuations, and the remaining terms arise from correlated offspring number fluctuations.

## S3 Evolutionary implications

To study the implications of decoupling noise, we consider our two-genotype model in the continuous time diffusion limit under constant selection (assuming small deviations in total population size),

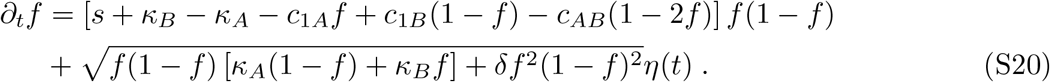

Where *η*(*t*) is standard gaussian white noise, ⟨*η*(*t*) ⟩ = 0 and ⟨*η*(*t*)*η*(*t*^*′*^) ⟩ = *δ*(*t* − *t*^*′*^). We use the approximate frequency mean and variance for the drift and diffusion terms from equations S12 and S10, respectively. We have combined several parameters for notational simplicity,

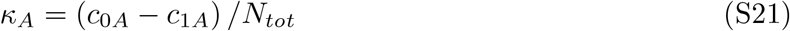

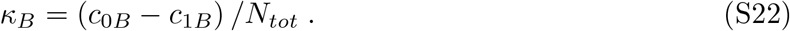

This model is similar to those analyzed in previous studies. Gillespie (1974)[8] studied the case where *δ* = 0 but *κ*_*A*_≠ *κ*_*B*_. Conversely, Melbinger and Vergassola (2015)[9] studied the case where *δ*≠ 0 but *κ*_*A*_ = *κ*_*B*_.

### S3.1 Fixation probability

Given that the genotype/mutant gets introduced into the population at some initial frequency *f*_0_, what is the probability that it will fix in the population? To find this, we’ll use the Kolmogorov Backward Equation (KBE) for this system. In general, the KBE is,

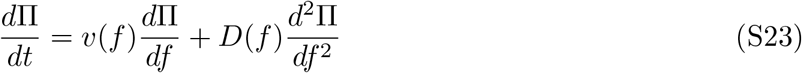

In this case, the drift and diffusion terms are given by,

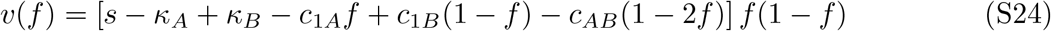

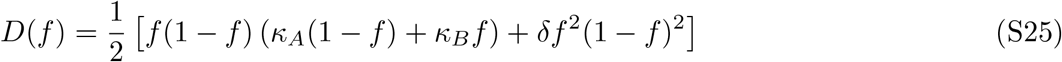

The steady state solution of equation S23 gives us the fraction of trajectories that end up at each of the absorbing states (here: 0 or 1). After setting equation S23 to 0, and applying the boundary conditions Π(0) = 0 and Π(1) = 1, we can rearrange and integrate to obtain a general solution for the fixation probability,

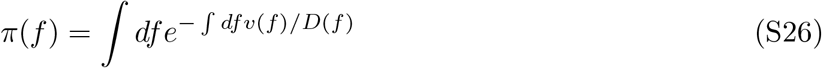

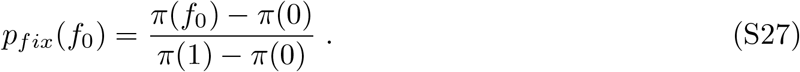

For our system, the fixation probability becomes,

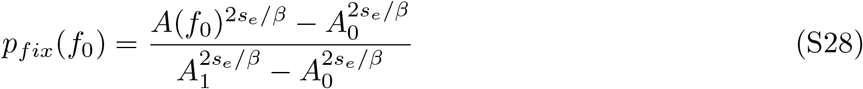

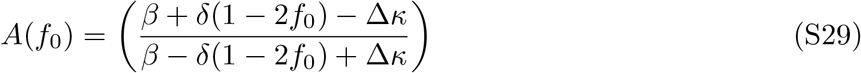

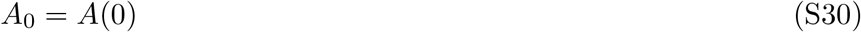

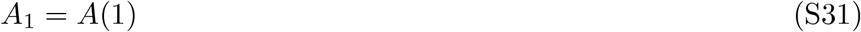

Or expanded,

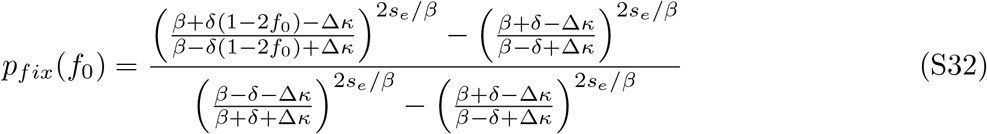

Which are in terms of new compound parameters,

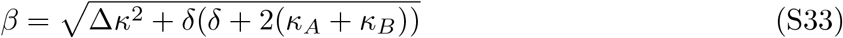

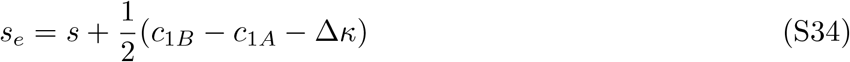

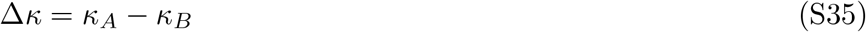

We are typically interested in the case where the initial frequency of the minor genotype is small, perhaps because it arose via spontaneous mutation in the population, or an individual with that genotype migrated to the population. Thus, we would like to focus on the limit of small initial frequency, *f*_0_ ≪ 1. We start by Taylor expanding equation S29 around *f*_0_ = 0,

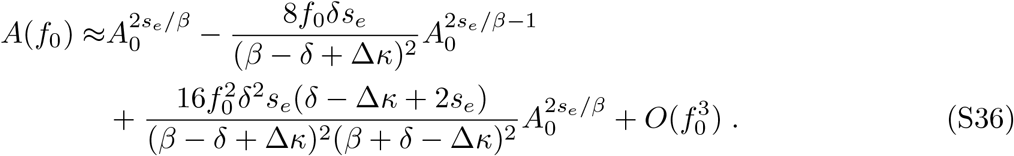

Following some algebra, we see that the second-order term in the expansion is negligible compared to the first-order term when,

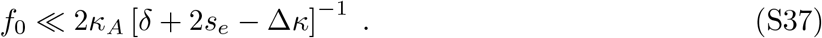

In this limit, we can truncate equation S36 to first order in *f*_0_; the fixation probability becomes,

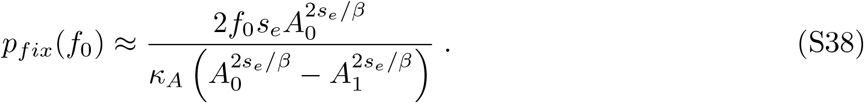

With this expression, we now wish to analyze the behavior of *p*_*fix*_ in the limit of weak selection, *s*_*e*_ → 0. We thus expand in *s*_*e*_,

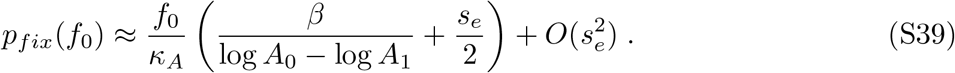

By comparing the first two terms, we see that *p*_*fix*_ is approximately constant at low *s*_*e*_,

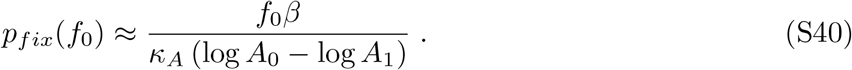

This approximation is valid when 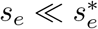, where,

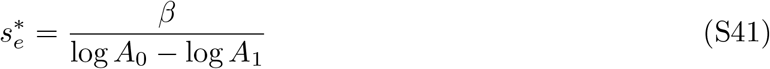

We recover the classical expression for the critical fitness effect when decoupling noise is weak, *δ* ≪ *κ*, and when Δ*κ* = 0,

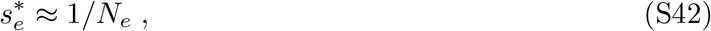

and thus we also see that the fixation probability when 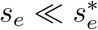is simply *f*_0_ in this case.

Now we turn to analyzing the opposite limit, where selection is strong. We will go back to the unapproximated expression for *p*_*fix*_ in equation S32. Rearranging, we obtain,

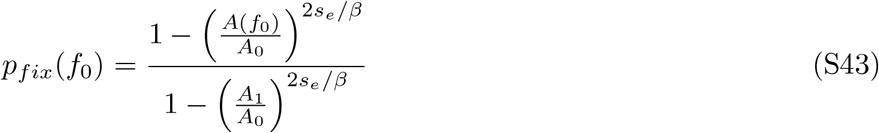

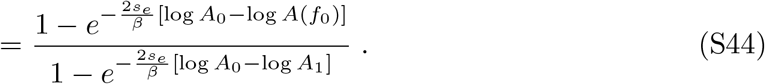

When 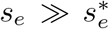and Δ*κ* = 0, we can expand in either small *δ* or small *N*_*e*_ to recover Kimura’s classical fixation probability [10],

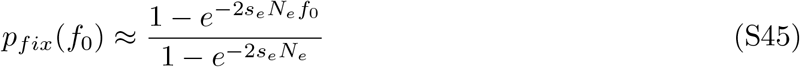

This approximation is also valid in the limit of weak decoupling noise, 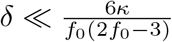.

### S3.2 Site frequency spectrum

The site frequency spectrum (SFS) describes the expected density of derived alleles at a given frequency; specifically *p*_*SF S*_(*f*)*df* is the number of derived alleles in the frequency range [*f*− *df /*2, *f* + *df /*2] [11]. We calculate the SFS for alleles affected by decoupling noise by leveraging previously described approaches [12, 13] along with our Langevin equation (equation S20). We find a general closed form solution for the SFS in terms of the frequency of alleles with parameters from *A*,

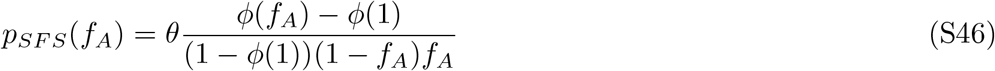

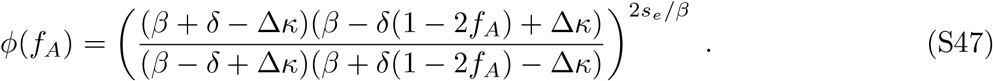

When *f* = 0.5 and Δ*κ* = 0, the SFS reduces to,

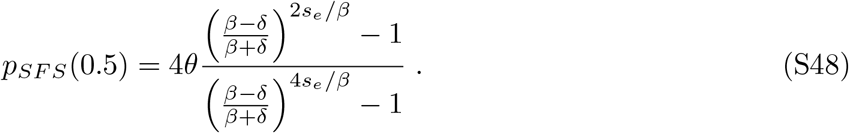

In the limit that *δ* ≫ 1*/N*_*e*_,

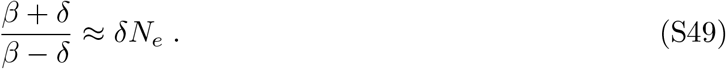

and,

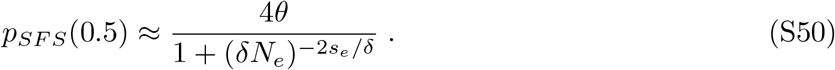

Similarly, the limit of the SFS as *f* → 1 is,

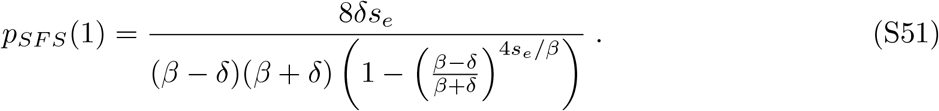

Applying the limit that *δ* ≫ 1*/N*_*e*_,

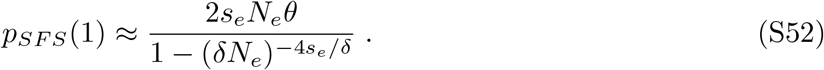

If we are interested in the weak selection limit, where *s* → 0, and again Δ*κ* = 0 then equation S46 becomes,

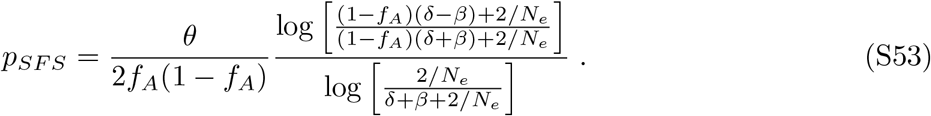

In the limit of strong decoupling noise, *δ* ≫ *N*_*e*_, equation S53 becomes,

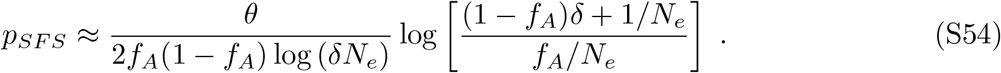

In the high frequency limit, (1 − *f*) ≪ 1,

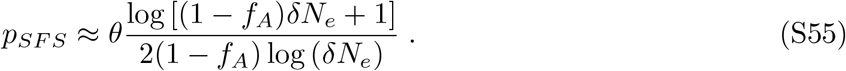

We see that when we additionally apply the limit that (1 ™ *f*_*A*_) ≫ (*δN*_*e*_)^−1^, we see that the SFS shows an uptick in the intermediate-high frequency regime,

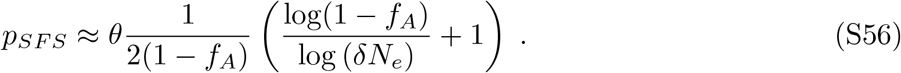

We find that in the limit of weak correlated fluctuations, *δ* ≪ 1*/N*_*e*_, and when *κ*_*A*_ = *κ*_*B*_, equation S46 reduces to the classical expression [14] for the site frequency spectrum for mutations under constant selection, i.e.

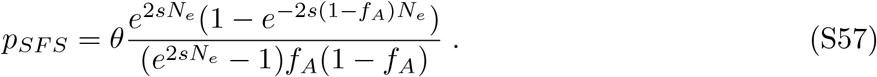

## S4 Chaotic dynamics

### S4.1 Variance between biological replicates

Here we show how exponentially increasing variance between biological replicates is equivalent to exponentially diverging trajectories, which is characteristic of chaotic dynamics. Specifically, in a one-dimensional system where we observe two trajectories, *f*_*i*_ and *f*_*j*_, initially separated by *ϵ*, we expect that at small times, their distance from each other will grow exponentially,

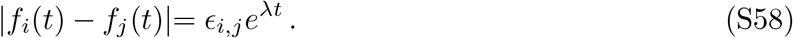

In chaotic systems, *λ >* 0. If we observe *N* initially nearby trajectories, labeled *f*_*i*_ (dropping the time index for simplicity), then the variance between trajectories is,

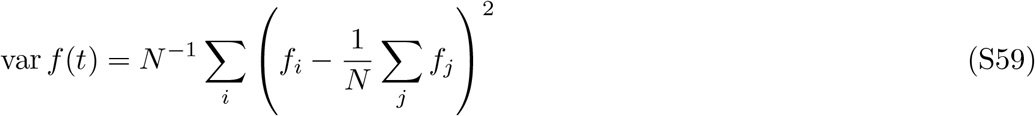

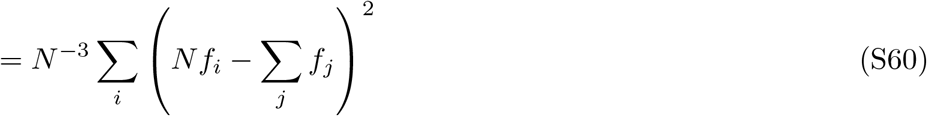

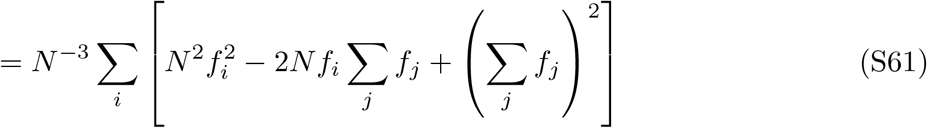

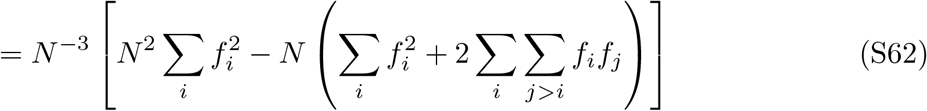

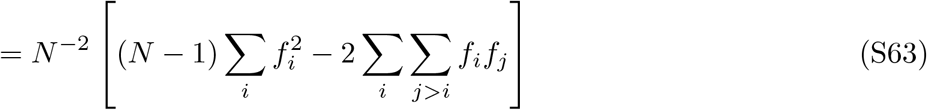

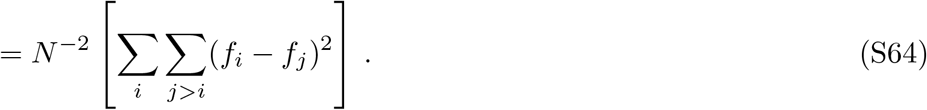

From equation S58, we can then replace (*f*_*i*_ − *f*_*j*_)^2^ with *ϵ*_*i,j*_*e*^2*λt*^, obtaining,

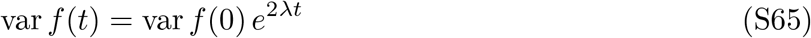

---

**Figure S1:**
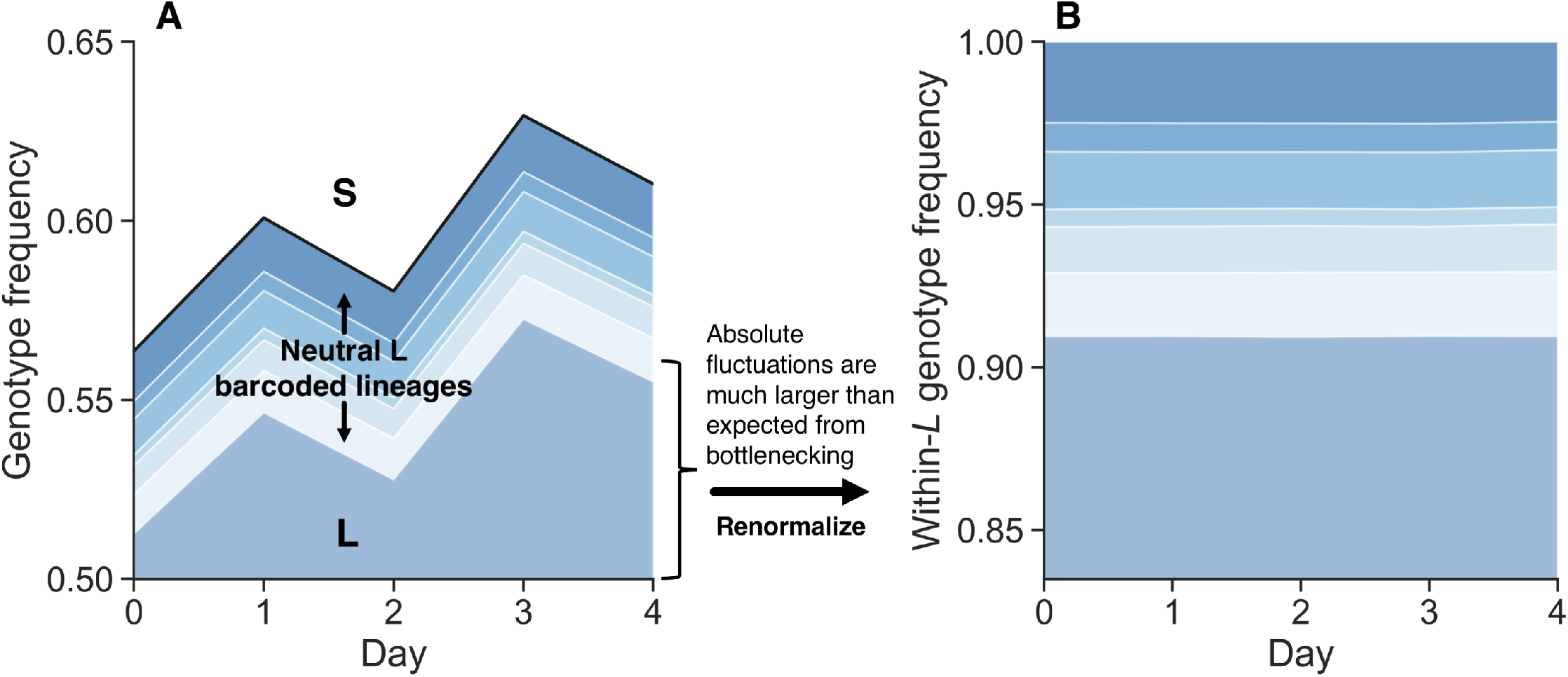
Fluctuations within the barcoded *L* library and relative to *S* (related to Figure 1A-B). (**A, B**) Randomly barcoded libraries of *E. coli* strains *S* and *L* were propagated together in their native serial dilution environment (previously reported data [15]). In the muller plot representation of lineage sizes, we see that the total frequency of *L* relative to *S* shows large fluctuations. However, neutral barcoded lineages within *L* show substantially smaller fluctuations relative to each other.

**Figure S2:**
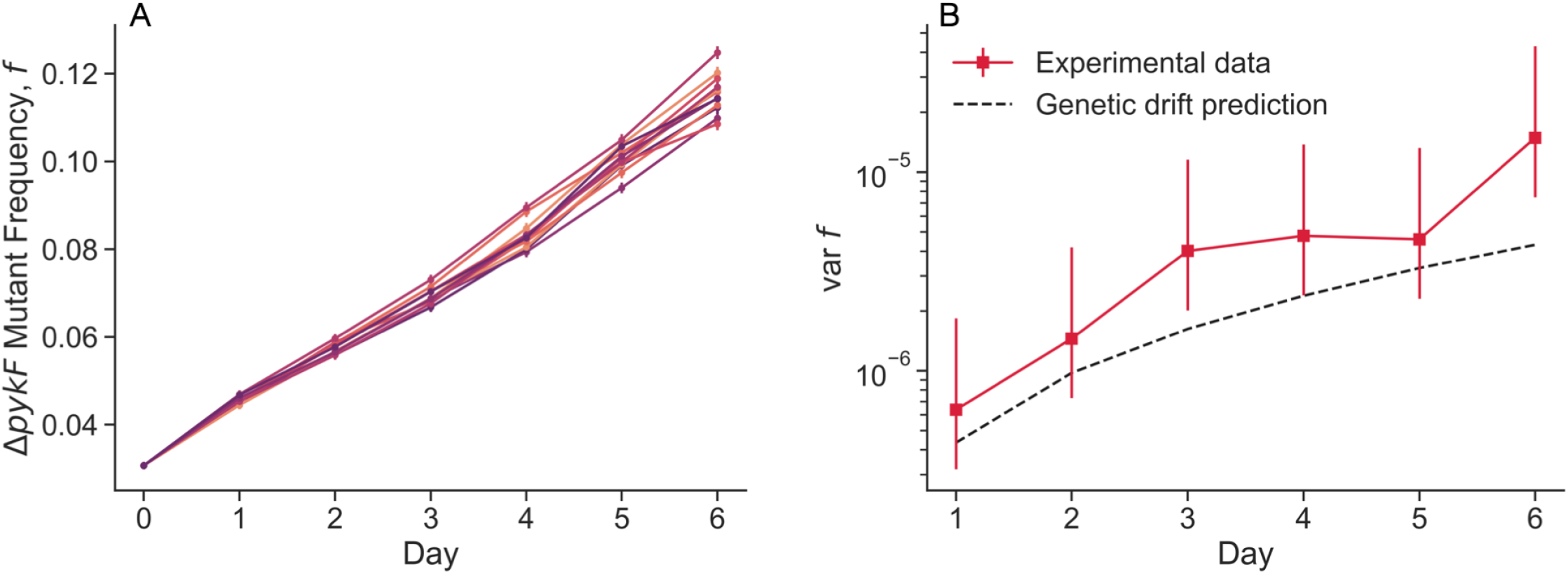
Coculture of REL606 and Δ*pykF* mutant (related to Figure 1D-E). (**A**) Cocultures were split at day 0 into 8 biological replicates, then propagated separately in the same environment. Flow cytometry measurements were taken every day. (**B**) We computed the variance between biological replicates, and compared it to the expectation that classical genetic drift is the sole source of population stochasticity, assuming *N*_*e*_ = 10^5^. Error bars represent 95% CIs.

**Figure S3:**
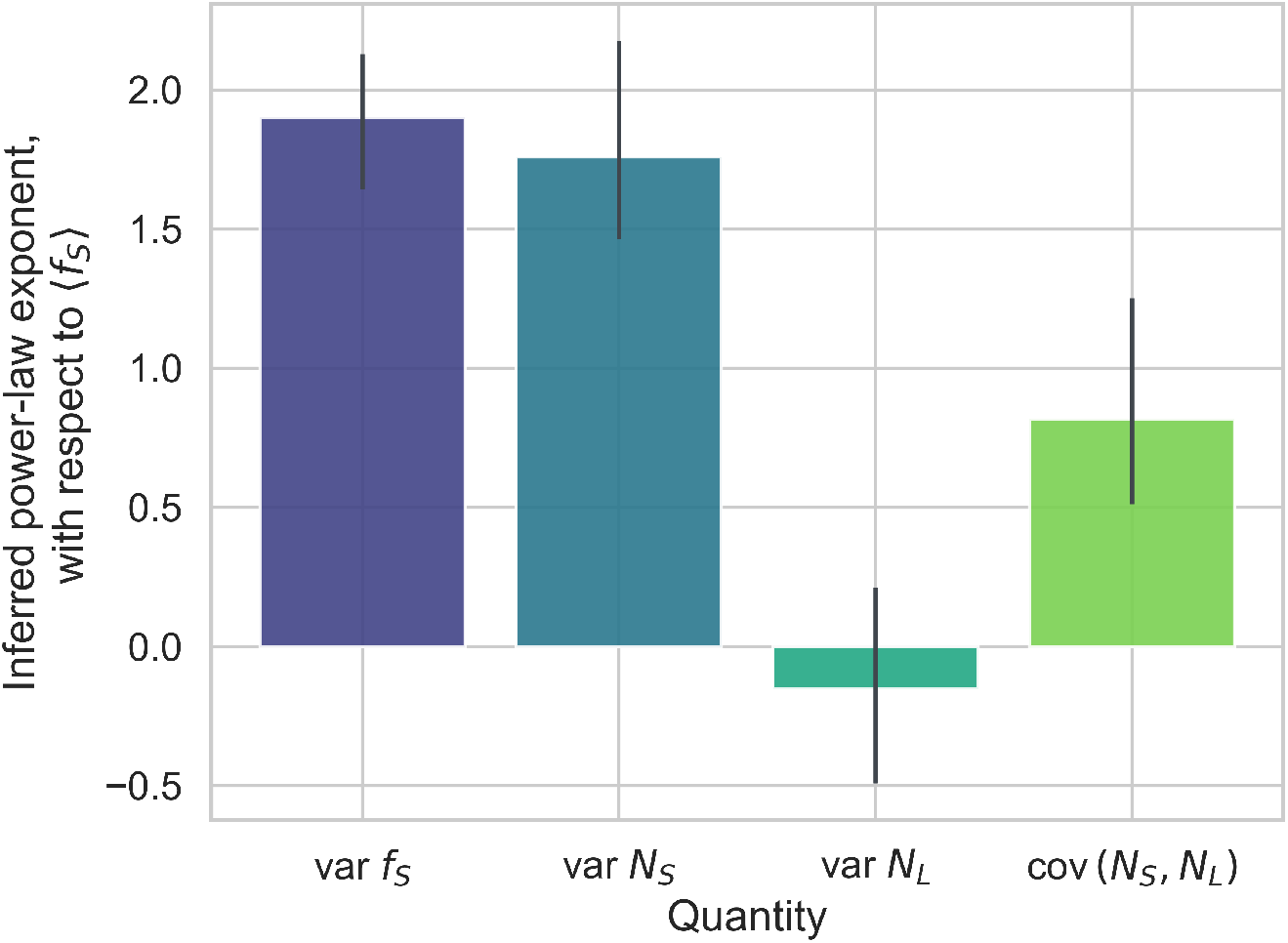
Inferred power-law exponents for variance and covariance quantities, as a function of mean *S* frequency (related to Figure 2). Error bars represent 95% confidence intervals, computed via bootstrapping.

**Figure S4:**
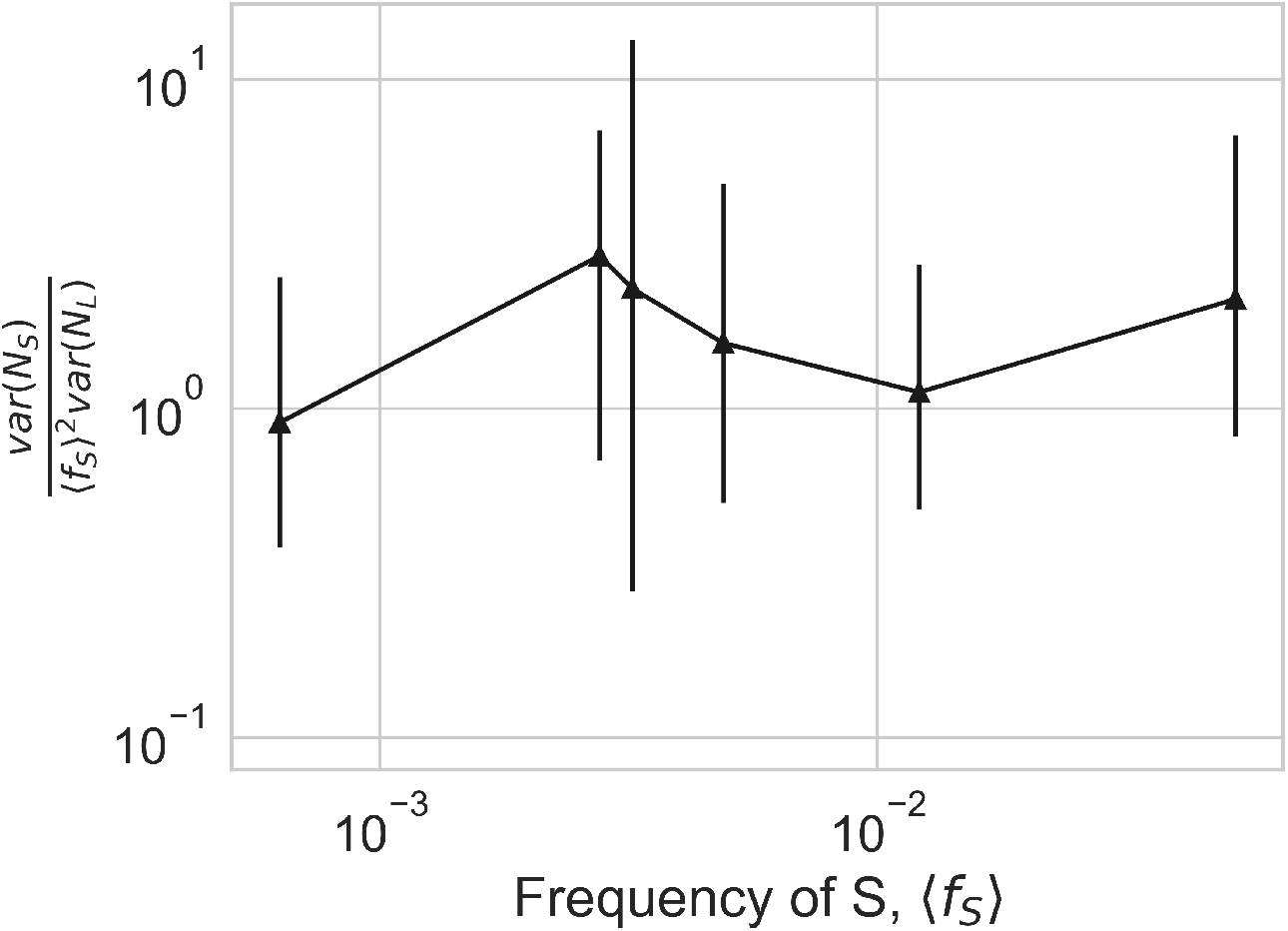
Ratio of abundance fluctuation strengths between *S* and *L*, including the *f* ^2^ factor for *S* (related to Figure 2). The ratio is equal to *c*_1*S*_*/c*_1*L*_ under our model. We see that the ratio is a small factor of order 1, indicating that *S* and *L* fluctuate by approximately the same amount. Error bars represent 95% confidence intervals.

**Figure S5:**
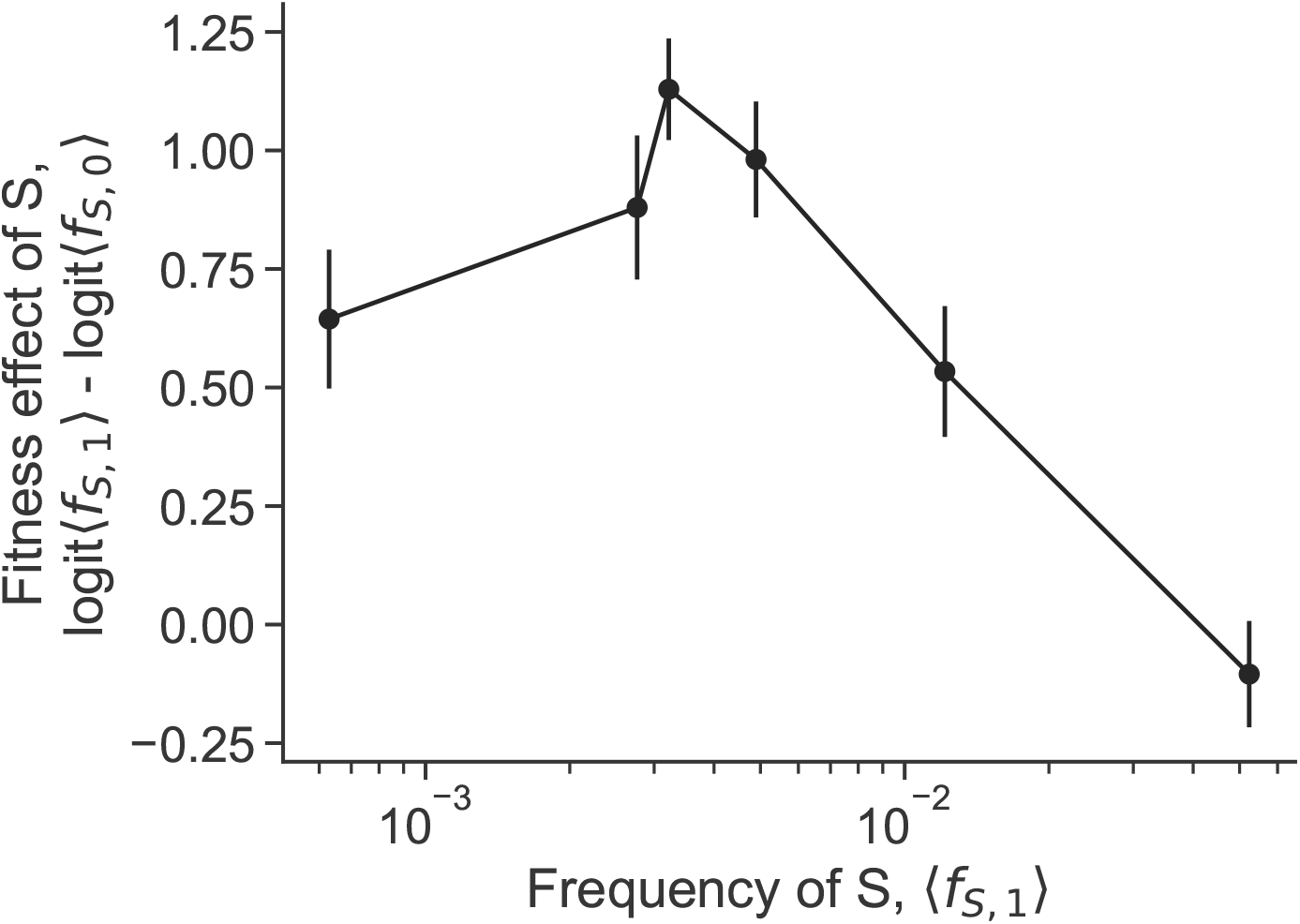
Frequency-dependent fitness effects of *S* and *L* (related to Figure 2). We computed the fitness effect of *S* relative to *L* across frequency. Consistent with prior measurements [16, 17], we find negative frequency-dependent fitness effects. Error bars represent 95% confidence intervals.

**Figure S6:**
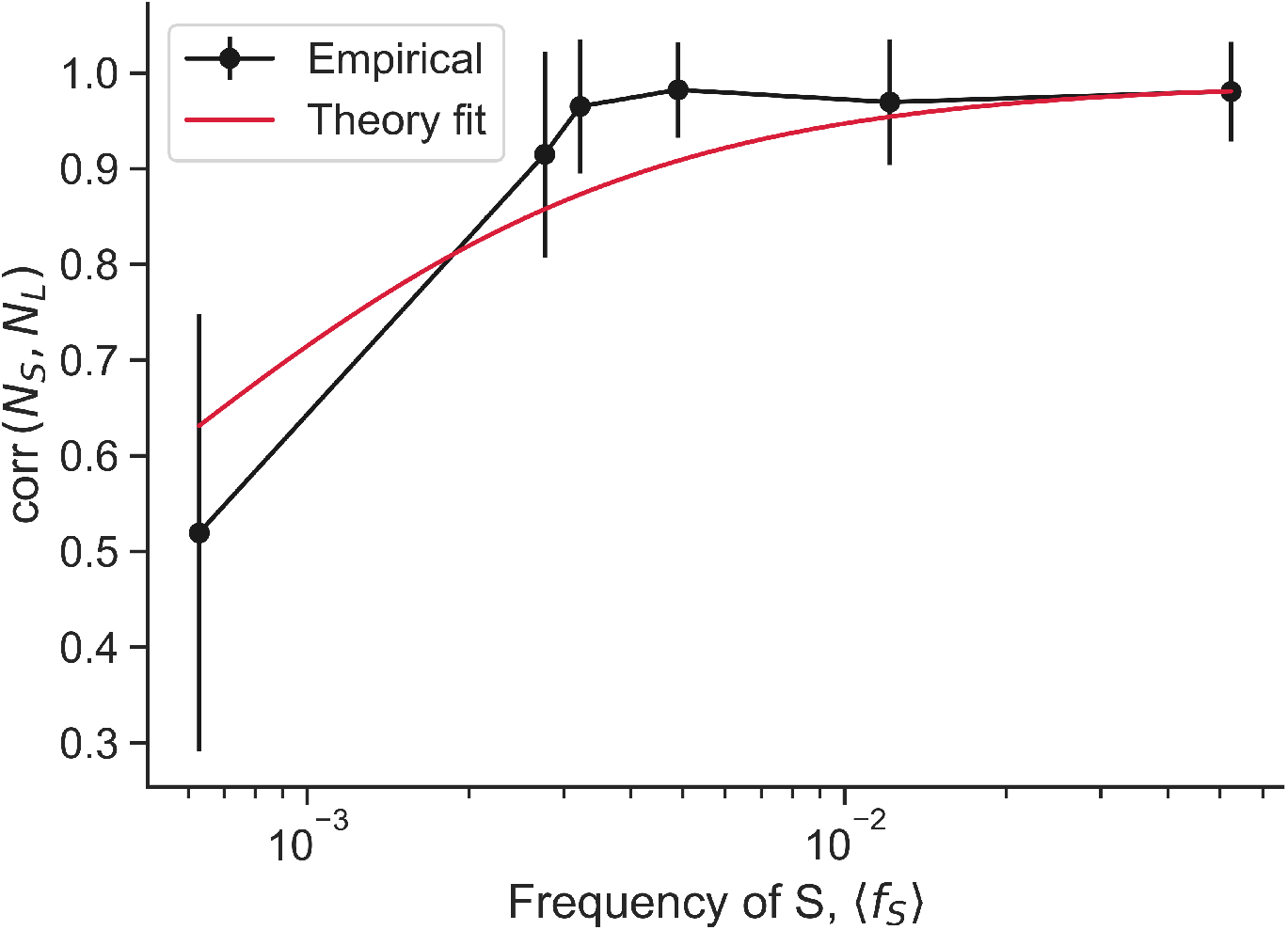
Correlation between the abundance of *S* and *L* (related to Figure 2). Our model predicts that the correlation between abundance should be constant when decoupling noise is dominant 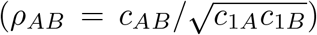. But once classical genetic drift becomes non-negligible, the correlation should decline; this perhaps indicates that the lowest data point is around the cross-over between the drift- and decoupling-dominated regimes. We fit the theoretical curve to our experimental data via ordinary least squares regression. Error bars are standard errors.

**Figure S7:**
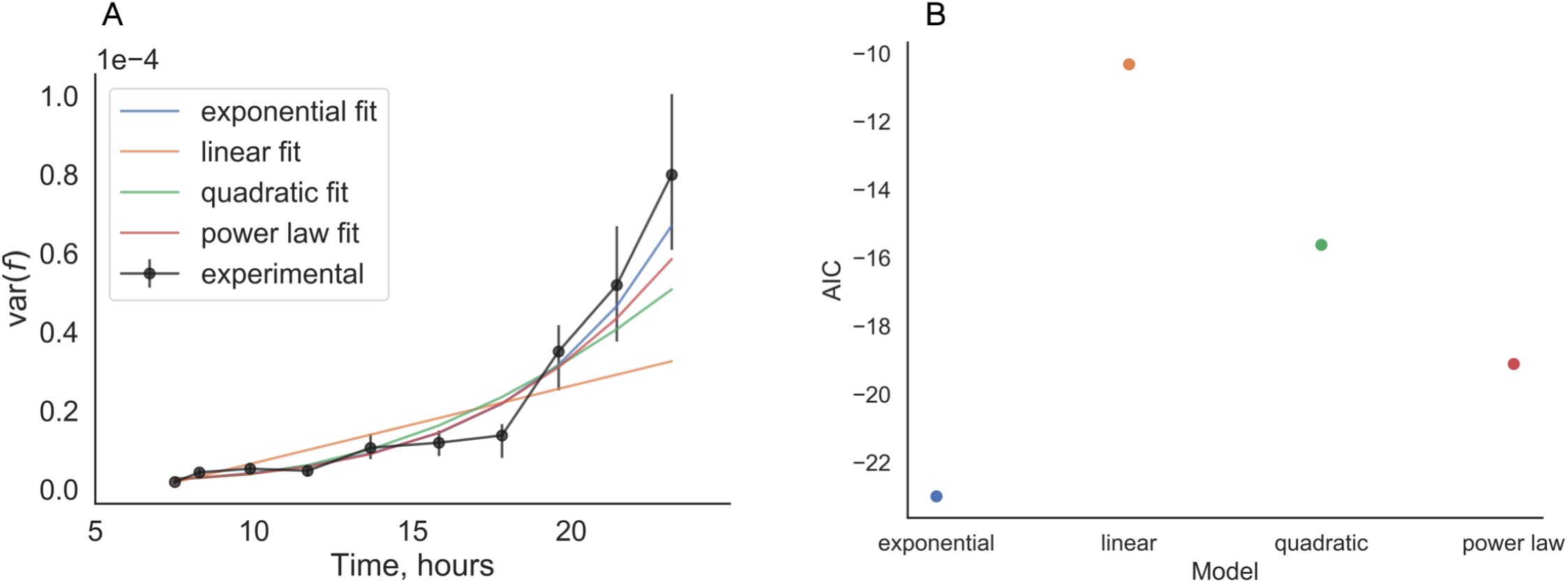
Comparison of curve fits to the variance of overnight frequency trajectories (related to Figure 4). (**A**) Best fits of various curves to the experimental variance measurements. (**B**) Akaike Information Criterion (AIC) of fitted models to experimental data, to evaluate goodness-of-fit. We computed FDR-corrected p-values to evaluate if the difference between the AIC of the exponential model compared to the other fits was signficant. The p-values were *p* =0.018, 0.058, and 0.022 for the comparison to the linear, quadratic, power law models respectively.

**Figure S8:**
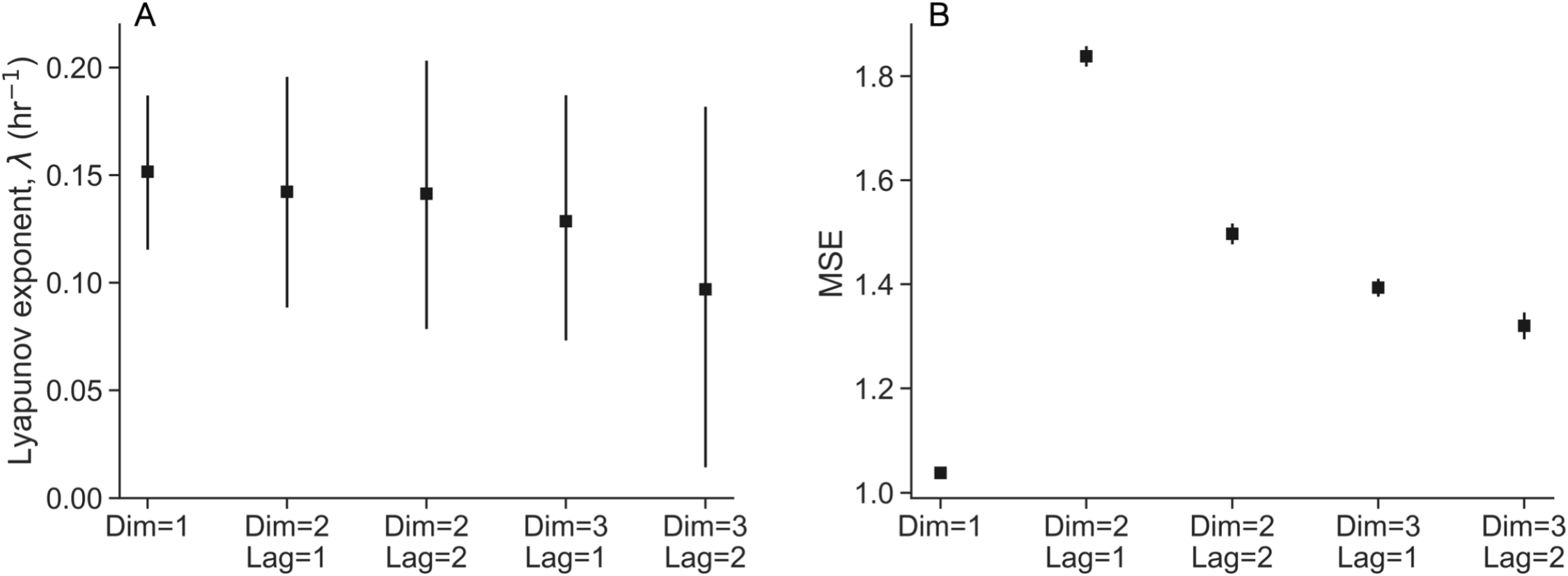
Cross-validation to compute Lyapunov exponent (related to Figure 4). We embedded the overnight trajectories in dimensions 1-3 by using a lag of 1-2 time points. (**A**) We computed the Lyapunov exponents using the method of Rosenstein et al. [18] for all conditions. (**B**) In order to choose which set of dimension and lag hyperparameters were most suitable for our data, we performed shuffle splitting cross-validation, and computed an out-of-sample mean squared error (MSE) for all conditions. All error bars represent 95% confidence intervals.

**Figure S9:**
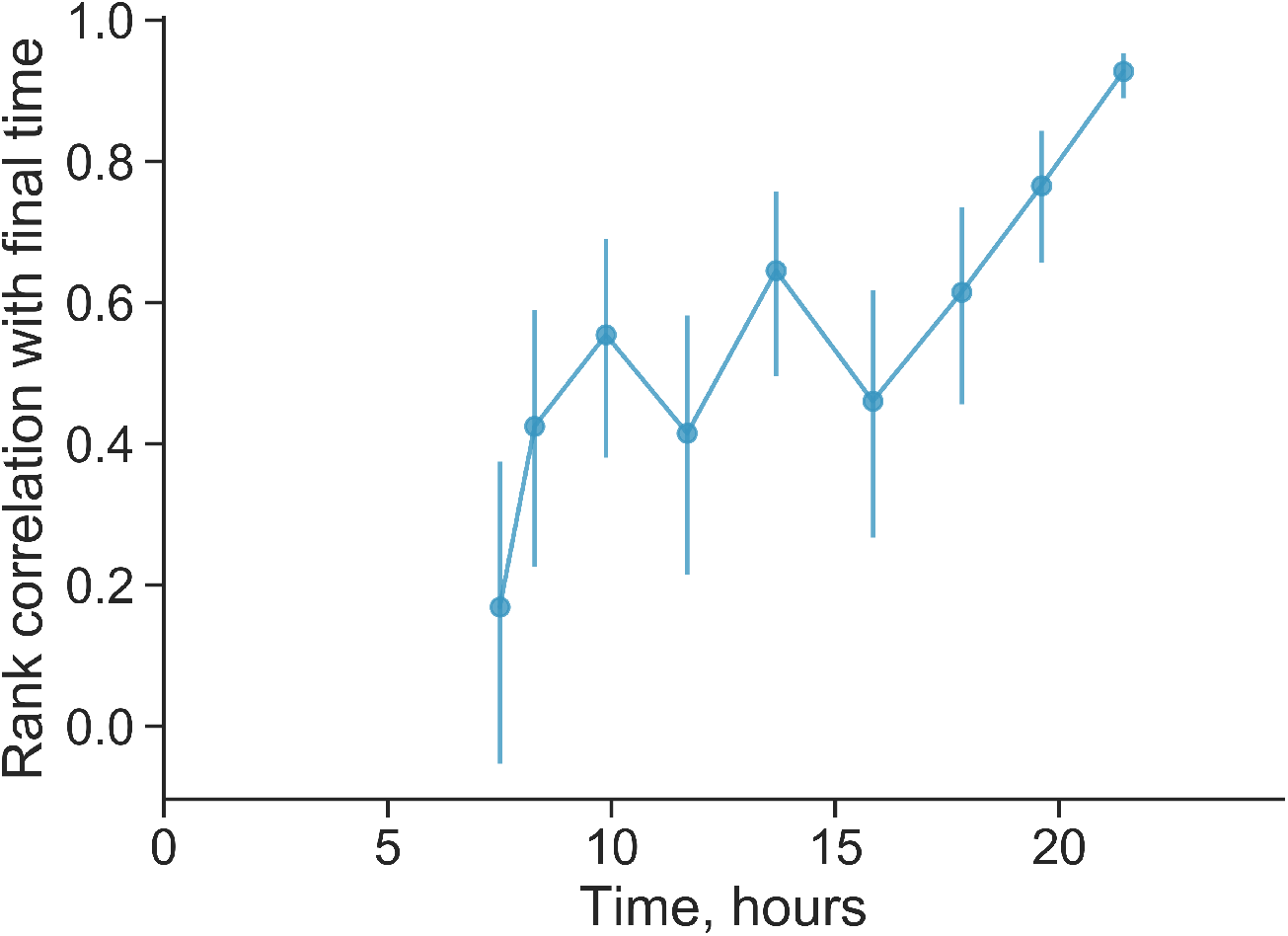
We quantified the rank correlation of replicates with its rank at the final time (related to Figure 4). The increase in correlation in the last 7 hours further shows how the decoupling noise accumulates over time, and that the rank position of each replicate is not “frozen in” earlier in the time course.

**Figure S10:**
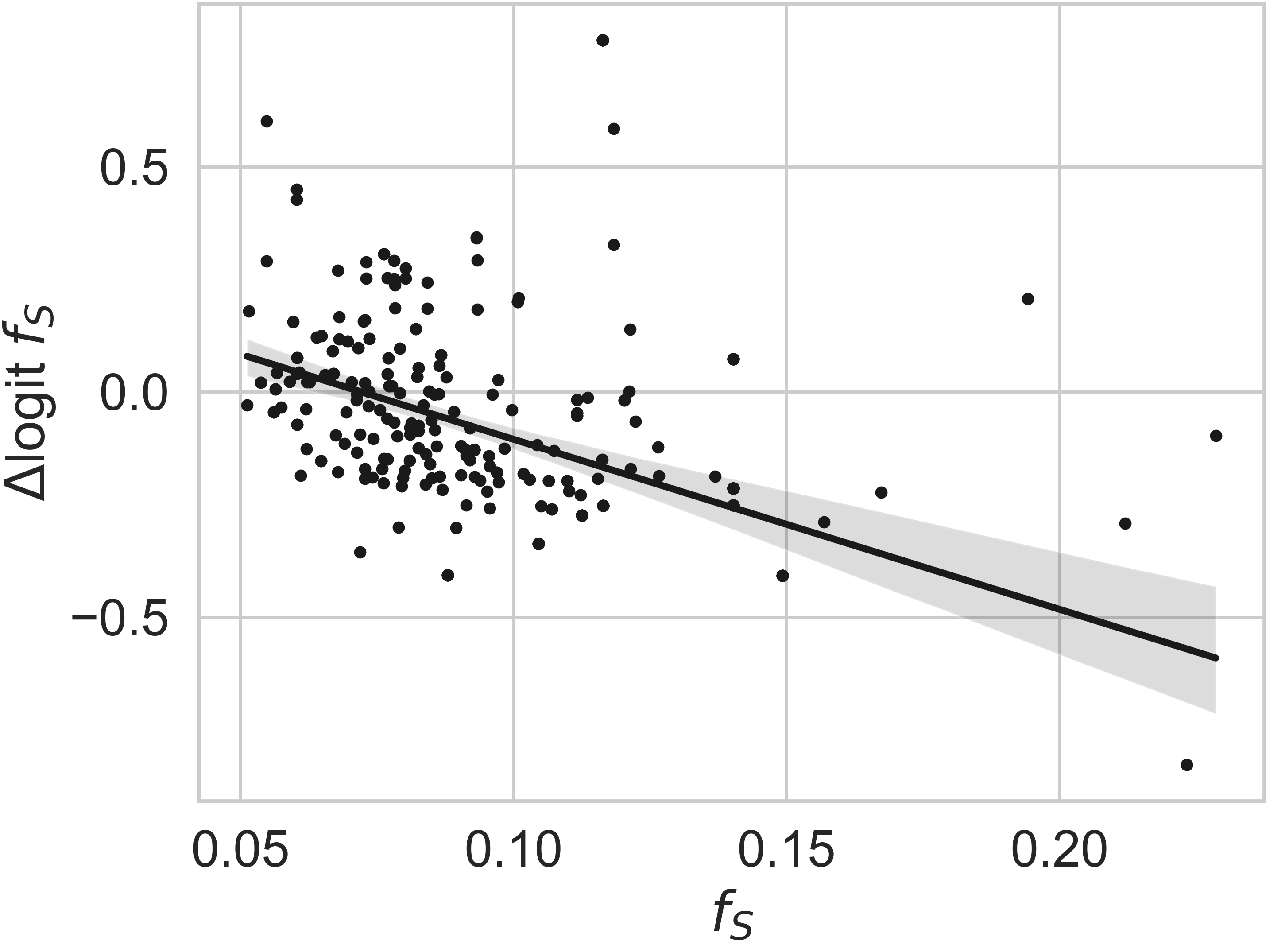
Frequency-dependent fitness effects of *S* and *L* (related to Figure 5). Scatter points represent change in logit frequency across a single day. Fitted line represents marginal frequency-dependent effect from the hierarchical model, described in the main text.

**Figure S11:**
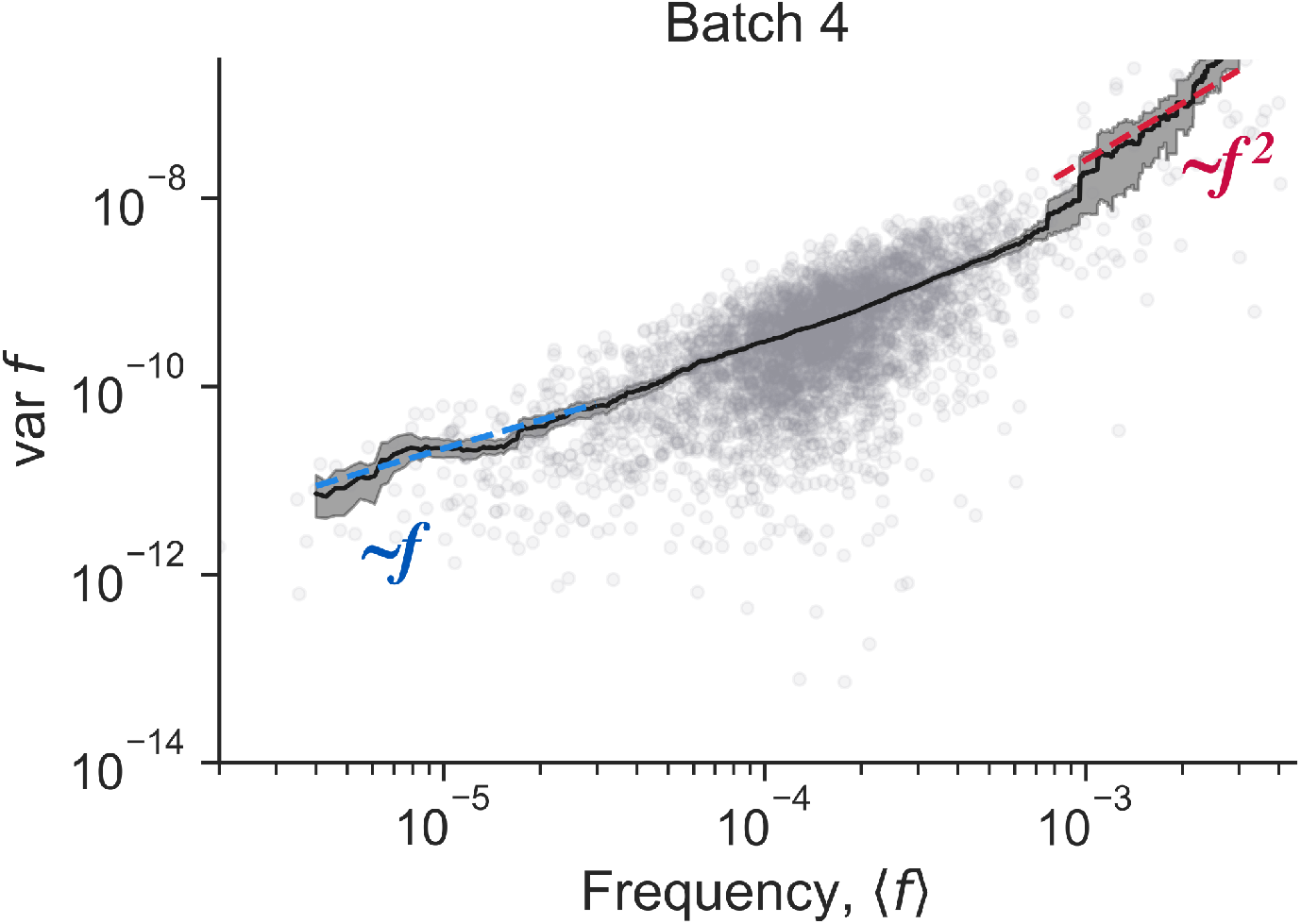
Frequency variance scaling in batch 4 in Venkataram et al.[19] data (related to Figure 6). Colored dashed lines represent the indicated scaling. Unconstrained power-law fits yields power law exponents of 2.15 ± 0.13 at high frequencies and 1.03 ± 0.08 at low frequencies.

**Figure S12:**
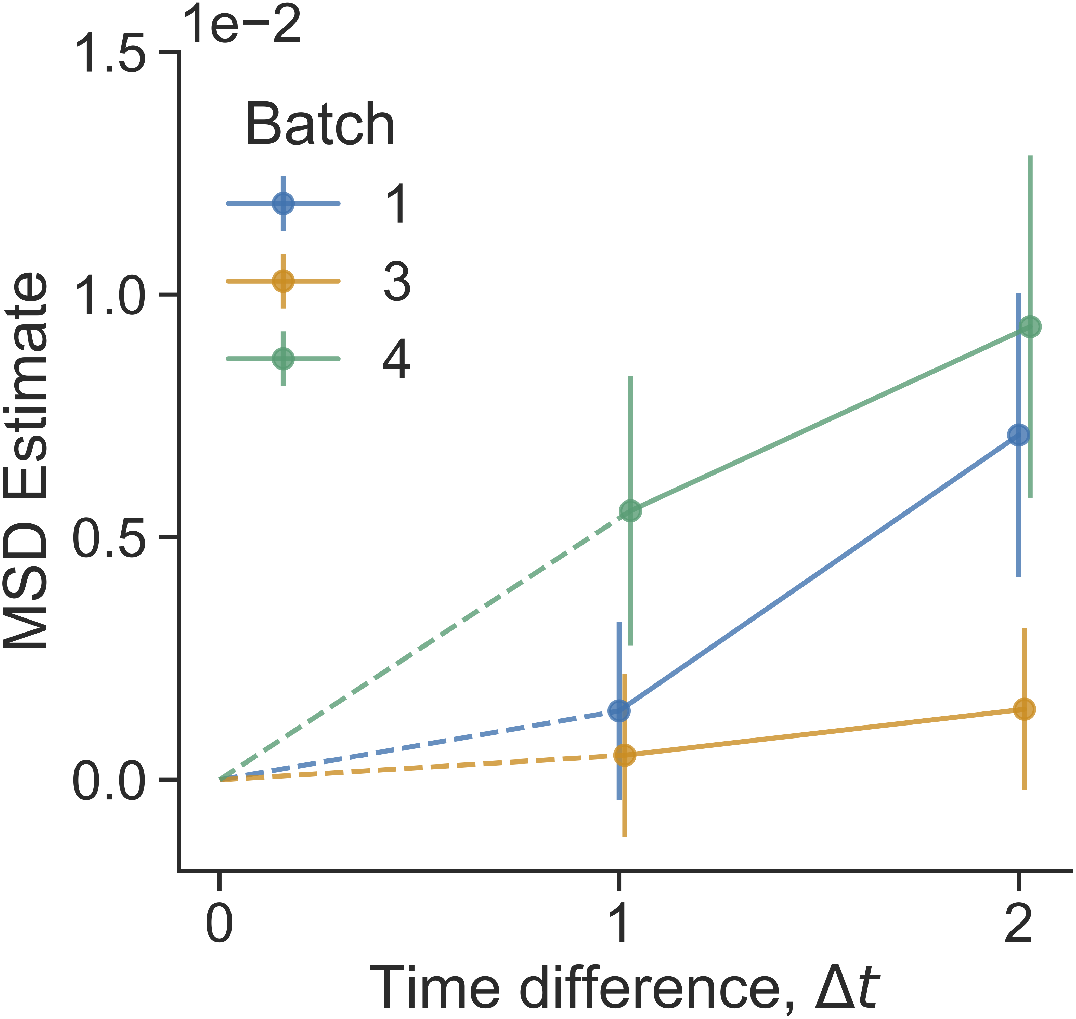
Mean squared displacement estimate computed with different frequency range, from Venkataram et al.[19] data (related to Figure 6B). Here, we focused on barcodes from a thinner mean frequency range (compared to the main text), 7 · 10^−4^ to 2 · 10^−3^. As there were fewer barcodes in this range, the estimates are noisier. Error bars represent 95% CIs.

**Figure S13:**
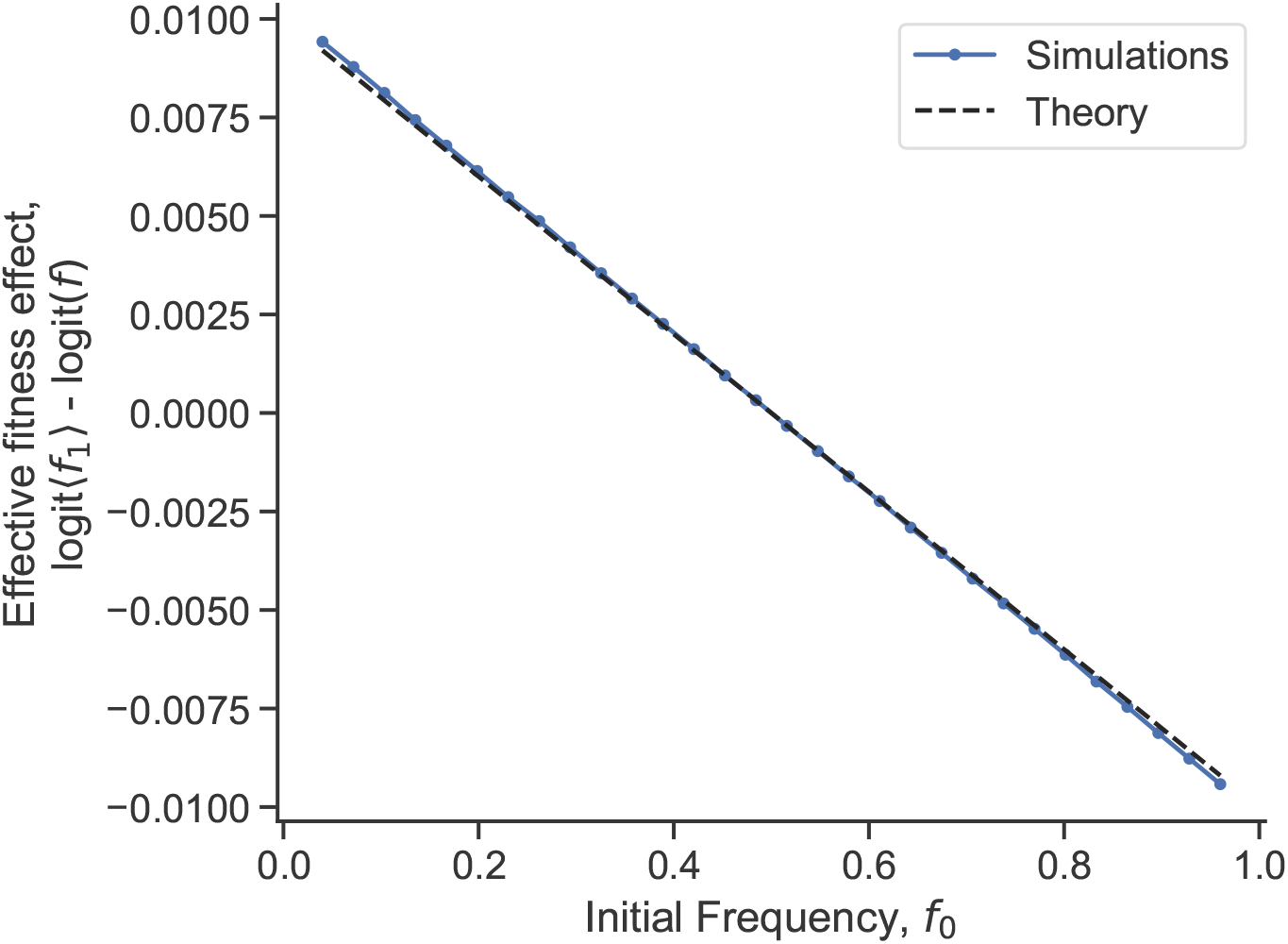
Effective frequency-dependent fitness effect induced by decoupling fluctuations (related to Figure 7). We simulated two uncorrelated population sizes with no selective difference, but that experience both genetic drift and decoupling noise, 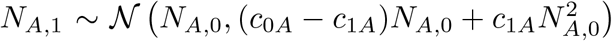 and 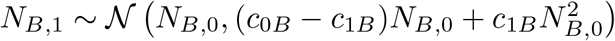. In the simulations, we used *c*_0*A*_ = *c*_0*B*_ = 1 and *c*_1*A*_ = *c*_1*B*_ = 0.01 (and *c*_*AB*_ = 0). We defined *f* = *N*_*A*_*/*(*N*_*A*_ + *N*_*B*_). We find that the stochastic decoupling noise induces an effective frequency-dependent fitness effect, which agrees well with the theoretical prediction, *c*_1*B*_(1 − *f*_0_) − *c*_1*A*_*f*_0_ − *c*_*AB*_(1 − 2*f*).

**Figure S14:**
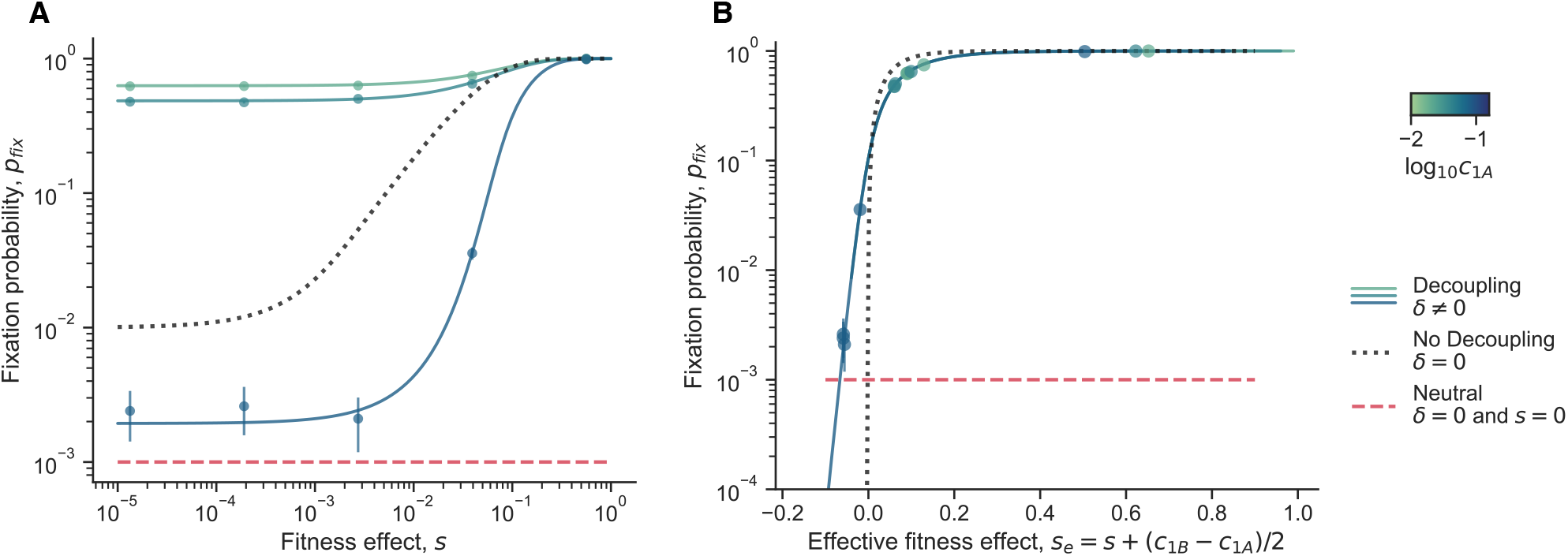
The dependence of *p*_*fix*_ on *s*_*e*_ (related to Figure 7). We show simulation results (round markers) and analytical predictions (solid lines) for the fixation probability, keeping *δ* constant but varying *c*_1*A*_. We consistently use *ρ* = 0, so that *c*_1*B*_ = *δ* − *c*_1*A*_. We show the fixation probability as a function of (**A**) the raw fitness effect, *s*, and (**B**) the effective (constant) fitness effect *s*_*e*_ = *s* + (*c*_1*B*_ − *c*_1*A*_)*/*2. When we consider the fixation probability as a function of *s*_*e*_, we see that the curves collapse into a single line. Additional parameter values used: *δ* = 0.2, *f*_0_ = 10^−2^, and *N*_*e*_ = 10^3^.

**Figure S15:**
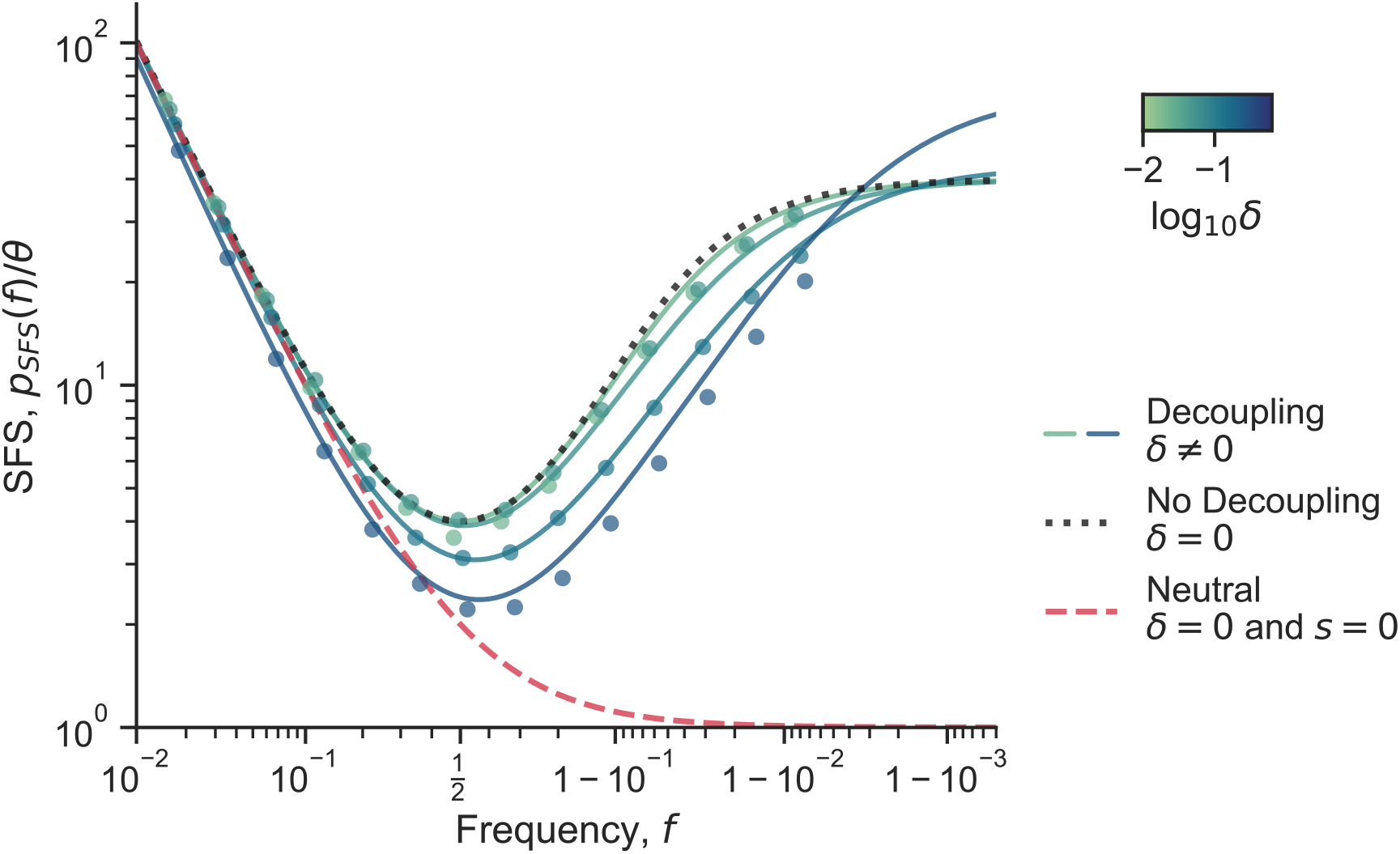
Site frequency spectrum with decoupling noise and constant selection. The same as Figure 7B, except where *s* = 0.02. The blue-green solid lines represent full analytical solutions for dynamics with decoupling noise, across different values of *δ*. The round markers show simulation results. The black dotted lines represent the case where there are no decoupling noise, but there is still (constant) natural selection. The red dashed lines represents the case where there is neither decoupling noise nor natural selection. We used *c*_1*A*_ = *c*_1*B*_, *ρ*_*AB*_ = 0.5, *N*_*e*_ = 10^3^ and *s* = 0.02.

